# Aβ Antibodies Target Not Only Amyloid Plaques But Also Distinct Brain Cells And Vessels

**DOI:** 10.1101/2025.09.24.678300

**Authors:** Gehua Wen, Nils Lindblom, Xiaoni Zhan, Megg Garcia-Ryde, Tomas Deierborg, Asgeir Kobro-Flatmoen, Gunnar K. Gouras

**Author notes:** Correspondence: Gehua Wen, Gunnar K. Gouras.

## Abstract

**Background:** Antibodies against amyloid-β (Aβ) are the only therapies capable of slowing cognitive decline and reducing Aβ plaque burden in Alzheimer’s disease (AD). Yet the precise sites where antibodies engage Aβ in the brain, and the mechanisms that drive Aβ clearance, are not fully defined. Defining the Aβ antibody engagement with Aβ in brain is essential to understand how immunotherapy can be beneficial for AD.

**Methods:** We administered the well-establsihed N-terminal Aβ antibody 6E10 via intrahippocampal (IH), cisterna magna (CM), or intraperitoneal (IP) injection in AD mouse models. N-terminal Aβ antibodies were seen as most effective in AD mouse models. Antibody 6E10 was not only assessed in Aβ plaques, but also examined in association with diverse brain cells and vascular compartments. Glymphatic dynamics were evaluated following Aβ antibody treatment.

**Results:** As expected, Aβ antibody 6E10 bound to Aβ plaques but remarkably also localized to vulnerable neurons, such as hippocampal CA1 pyramidal cells, as well as microglia, astrocytes, oligodendrocytes, perivascular macrophages (PVMs), and blood vessels. Glymphatic function showed no significant alterations after antibody administration.

**Conclusions:** Aβ antibodies distribute not only to amyloid plaques but also to neurons, glial cells, and blood vessels. This study provides detailed localizations of antibody in the AD brain, offering valuable insights into cellular targets and spatial dynamics of Aβ antibody-based immunotherapy.

## Background

Aβ immunotherapy is the first therapy targeting underlying disease mechanisms to show cognitive benefits in clinical trials of patients afflicted by AD[1]. The mechanism generally viewed as most important to these beneficial, albeit still modest, benefits is antibody-mediated degradation of amyloid plaques by microglia. However, amyloid plaques have long been known not to correlate well with cognition in AD, and increasingly, more soluble forms of Aβ ‘oligomers’ and ‘protofibrils’ have been viewed as the more important pathogenic form(s) of Aβ[2]. For example, an inducible AD mouse model with plaques showed that turning off the AD mutant APP transgene rapidly improved behavior while plaques remained unchanged[3]. Further, reduced brain activity via sleep induction or unilateral whisker removal reduced amyloid plaques but increased synapse damage in AD mice[4]. Thus, there is a disconnect between Aβ induced synapse damage and amyloid plaques. Although it is still widely viewed that secreted extracellular Aβ initiates plaque formation[5] and that then amyloid plaques cause damage to surrounding synapses and neurons from release of pathological forms of Aβ[6], a growing body of evidence over the past few decades, starting with our study, has shown that Aβ begins to aberrantly accumulate and pathologically aggregate within AD vulnerable neurons prior to the formation of extracellular plaques. This was initially shown in human Down syndrome (DS)[7–9] and AD brains[7, 10–12], but then was even more extensively studied in numerous mouse and rat models of AD[13–17]. Clues that amyloid plaque removal may not be the only target for improvement of cognition with Aβ immunotherapy are studies that have shown beneficial effects of Aβ antibody F_ab_ fragments in AD model mice[18] and that AD mice crossed with Fc𝛄 receptor knockout mice still showed benefits from Aβ immunotherapy[19]. Moreover, behavioral improvement was reported in AD mice with an Aβ-targeting antibody that did not reduce amyloid plaques[20]. Additionally, behavioral impairment was reported prior to amyloid plaques in the 3×Tg-AD mouse, with improvement in behavior from Aβ immunotherapy upon reduction of the intraneuronal pool of Aβ prior to the formation of plaques[21]. Neuropathological evaluation of patients in the initial, but prematurely aborted, active human immunotherapy trial showed plaque removal but continued cognitive decline in patients who came to autopsy[22]. Thus, multiple lines of evidence support that Fc receptor mediated microglia degradation of amyloid plaques does not appear to be the only route to cognitive improvement in AD. Yet, based on clinical evidence that plaque removal correlates with success of therapeutic Aβ antibodies, the field currently emphasizes plaque reduction as the critical target[23].

What other mechanisms might be important for the protective effects of Aβ antibodies? It is possible that soluble extracellular Aβ oligomers are sequestered by Aβ antibodies, although there is limited experimental evidence for this [24]; such studies that experimentally induce elevated extracellular Aβ are also inconsistent with what one sees in the human extracellular fluid cerebrospinal fluid (CSF) with AD. In fact, the earliest validated biomarker indicative of AD pathogenesis is a drop rather than an increase in extracellular Aβ42 in CSF[25, 26]. In contrast to the dearth of evidence for early increases in a more soluble extracellular pool of Aβ42, there are now hundreds of articles reporting on the early intraneuronal accumulation of Aβ in human AD, DS and numerous rodent models of AD. Further, intraneuronal Aβ42 was shown by immuno-eletron microscopy to pathologically aggregate within dystrophic neurites prior to the appearance of plaques[16].

Aβ antibodies might protect against intraneuronal oligomeric Aβ. In a cellular study, we had previously provided mechanistic insights into cellular internalization of Aβ antibodies reducing intraneuronal Aβ and protecting against AD-like synapse alterations in cultured neurons from AD transgenic compared to wild-type mice[27]. While deciphering the precise cellular mechansims whereby this occurs is challenging to fully determine, overall the evidence pointed to induced degradation of intraneuronal Aβ by antibody. However such mechanistic cellular studies, which are again ongoing in our lab, very much require in vivo support, but up to now Aβ antibodies have not been shown to localize to neurons in brain.

Previous studies have indicated that the clearance of Aβ from the brain is disrupted in AD, a process that has been connected to glymphatic system dysfunction[28–32]. The glymphatic system, a perivascular pathway shunting interstitial fluids alongside arteries and veins, plays a key role in clearing metabolic waste from the brain. Its function is influenced by changes in the polarization of astrocytic aquaporin-4 (AQP4) around blood vessels[33, 34] and by perivascular macrophages[35, 36]. However, the effects of Aβ immunotherapy on glymphatic system function remain poorly understood.

Among passive immunotherapies targeting Aβ, multiple monoclonal antibodies (mAbs) with good tolerability and target specificity have been developed, yet early clinical trials mostly produced disappointing outcomes. Extensive preclinical work on Aβ immunotherapy in AD mice had empirically demonstrated the therapeutic particularity of antibodies directed against epitopes located within the N-terminal region of the Aβ peptide[37]. Aducanumab was the first potentially disease-modifying therapy that received accelerated approval from the US FDA, while its clinical efficacy remained under debate among experts[38, 39]. In contrast, mAbs targeting the central domain or C-terminal region of Aβ generally demonstrate poor clinical efficacy[40–42]. Wilcock et al. employed antibody 6E10 in an acute intracranial injection paradigm, revealing dynamic crosstalk between microglia and other immune cells and highlighting early immune communication events during Aβ immunotherapy[43].

In this study, we set out to more precisely identify the cellular targets in the brain of the widely used N-terminal Aβ IgG1 monoclonal antibody 6E10. Prior studies had shown that Aβ antibodies bind to amyloid plaques when either intracranially or intravenously injected into AD transgenic mice[44, 45], but did not further investigate additional localizations of antibodies. Here, we show that Aβ antibodies also selectively localize to specific cell populations and blood vessels in the brain of AD mouse models. Intraneuronal Aβ42 accumulation represents a selective cellular vulnerability in AD, leading to synaptic dysfunction and neurotoxicity[46]. Remarkably, here we show Aβ antibody within AD vulnerable neurons, with a concomitant reduction in intraneuronal Aβ. Further, we show Aβ antibodies in microglia, perivascular macrophages, and astrocyte endfeet. Astrocytes are essential for brain function and Aβ clearance in AD[47]. Astrocytes[48] and perivascular macrophages[35] are also important for the function of the glymphatic system. Microglia are increasingly linked to AD, and Aβ has been shown to accumulate in microglia after immunotherapy[22]. Our observation that Aβ antibodies are present within microglia is of considerable interest and might be important to the effects of Aβ immunotherapy, and could affect microglia phenotype, but has so far not been explored. Perivascular macrophages are also increasingly studied in AD, and vascular amyloid is considered central to the major side effect associated with Aβ immunotherapy, namely amyloid-related imaging abnormalities (ARIA)[49]. The presence of Aβ antibodies within perivascular macrophages and blood vessels may contribute to the development of ARIA and/or influence their ability to clear Aβ from the brain.

## Results

### Unilateral hippocampal injection of Aβ antibody reduces plaques, reveals antibody within neurons, and activates astrocytes and microglia in 5xFAD mice

Prior studies had shown that N-terminal Aβ antibodies (e.g. antibody 3D6; humanized name: bapineuzumab) were particularly effective in reducing amyloid plaques in AD mouse brains[37, 50]. We first unilaterally injected either unconjugated (Fig. S1a, b) or Alexa Fluor 488-conjugated N-terminal Aβ antibody 6E10 into the hippocampus (IH) of 5xFAD mice; 24 or 72 h following injection, we analyzed the brains for antibody distribution and effects on plaques, astrocytes, and microglia (Fig. 1a; Fig. S1c, d). The unconjugated and fluorescently conjugated antibodies showed a similar pattern of labelling. However, since the signal of the conjugated antibody was considerably stronger, we proceeded with using this antibody for further experiments. Conjugated Aβ antibody distribution after IH injection showed prominent labeling in the injected hippocampus after 24 h, while after 72 h, the labeling was comparatively less pronounced (Fig. 1b, c, d). Remarkably, after 24 h, we detected a clear signal inside neurons, particularly in hippocampal field CA1 and the subiculum (Fig. 1e, f, g). MOAB-2 antibody was utilized to detect intracellular Aβ as well as plaques[51]. Labeling the same sections with anti-Aβ antibody MOAB-2 revealed a substantial degree of overlap with the intracellular labelling of using the 6E10 antibody, providing evidence that the intracellular target is indeed Aβ (Fig. 1h, i, j). In line with previous studies using Aβ antibodies[44], the 6E10 antibody was also detected in amyloid plaques (Figure 1k, l). Although no reduction (*p*=0.9762) in Aβ plaques from the Aβ antibody was observed in the hippocampus at 24 h post-injection (Fig. 1k, q), a significant reduction (*p*=0.0356) was detected at 72 h on the injected compared to the uninjected side (Fig. 1l, q). Similarly, at the 72 h but not 24 h time point, intracellular Aβ in CA1 neurons on the injected side was significantly reduced (uninjected-72 h vs injected-72 h, *p*=0.0012) compared to the uninjected side (Fig. S2a, b, c, d, e, f, i). Astrogliosis was assessed by measuring the expression of glial fibrillary acidic protein (GFAP), a marker of astrocytes. Microglial activation was evaluated by quantifying the expression levels of ionized calcium-binding adapter molecule 1 (Iba1), a microglia-specific marker. We observed increased activation of astrocytes (Fig. 1m, n, r) (uninjected-24 h vs injected-24 h, *p*=0.0014. uninjected-72 h vs injected-72 h, *p*<0.0001) and microglia (Fig. 1o, p, s) (uninjected-24h vs injected-24 h, *p*<0.0001. uninjected-72 h vs injected-72 h, *p*=0.002) on the injected side. However, the glial activation may be a result of the injected antibodies themselves and/or a consequence of the tissue damage from the injection itself. To better ascertain this, we performed unilateral IH injections of a fluorescent secondary antibody, which showed a significantly different pattern compared to that of the injected Aβ antibody (Fig. S1e). The secondary antibody was found predominantly at the surface of small blood vessels and did not visibly bind to plaques or label hippocampal neurons. Still, unilateral IH injection of the secondary antibody induced widespread activation of astrocytes and microglia in the hippocampus to a similar extent as that observed following the Aβ antibody injection (data not shown).

**Fig. 1.**
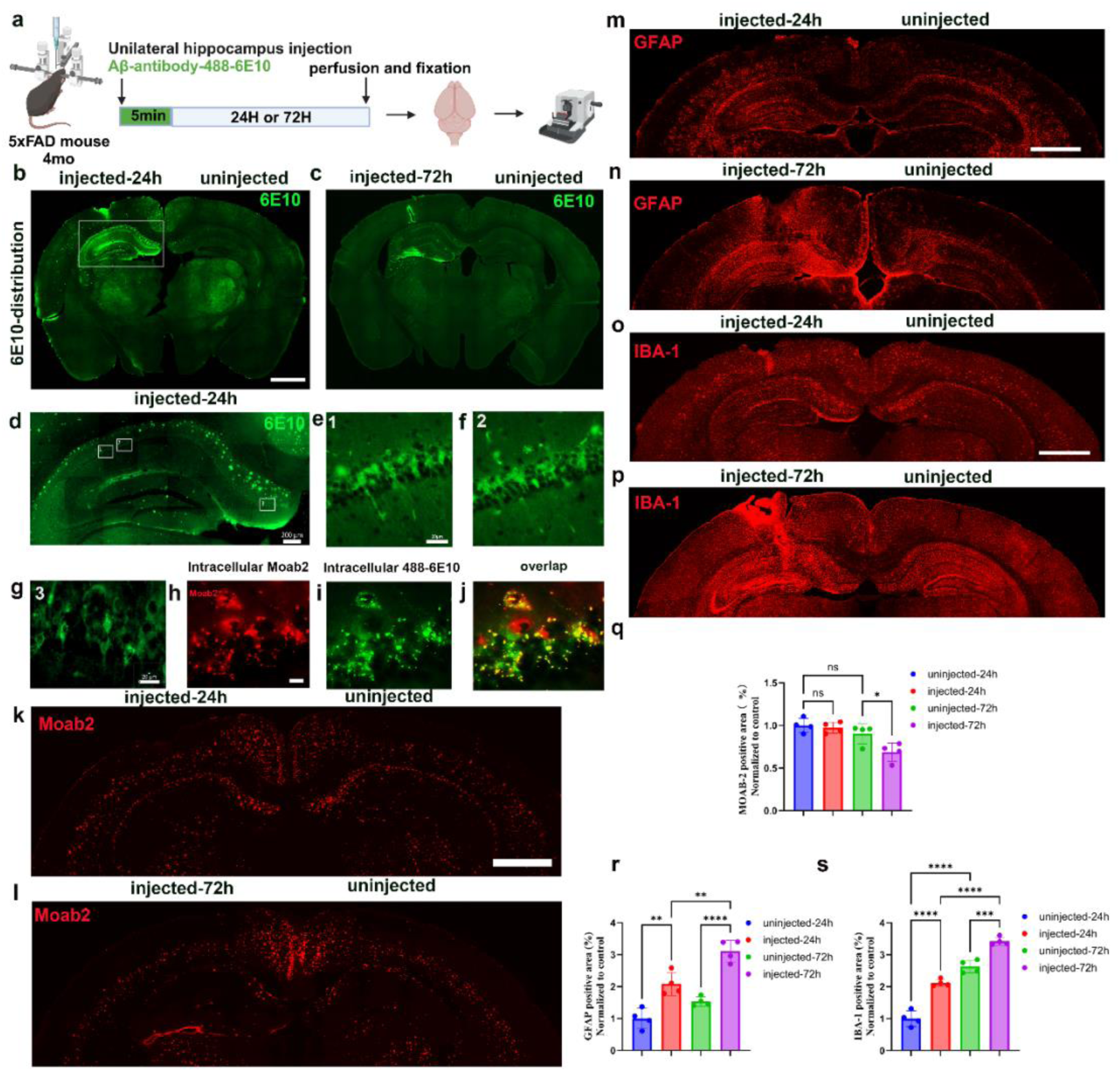
Unilateral intrahippocampal (IH) injection of Aβ antibody 6E10 is detected in neurons, reduces hippocampal plaques, and activates astrocytes and microglia in 5xFAD mice. (a) 5 μl of conjugated 6E10 antibody was injected into one side of the hippocampus of 4-month-old 5xFAD mice. (b, c) Representative images of Aβ antibody 6E10 24 and 72 h after IH injection show elevated antibody on the injected compared to uninjected side, as well as strong antibody labelling of plaques; scale bar = 1 mm. (d) At 24 hours post-injection, Aβ antibody 6E10 distributions in the injected hippocampus. Three regions in CA1 and subiculum were used for analysis; scale bar = 200 μm. (e, f, g) The three selected areas in Fig. 1d showed intracellular distributions of the Aβ antibody in neurons, scale bar = 20 μm. (h) Antibody MOAB-2 labeling in the CA1 for intracellular Aβ indicates that the intracellular Aβ antibody (i) colocalizes with intracellular Aβ (j), scale bar = 10 μm. The MOAB-2 levels in the hippocampus at 24 h (k) and 72 h (l) after unilateral IH injection; scale bar = 1 mm. GFAP levels in the hippocampus at 24 h (m) and 72 h (n) after Aβ antibody injection; scale bar = 1 mm. IBA-1 levels in the hippocampus at 24 (o) and 72 h (p) after Aβ antibody injection, scale bar = 1 mm. Quantification of MOAB-2 (q), GFAP (r), and IBA-1 (s) labeling in the hippocampus. N=4 per group. Data are represented as mean ± SD. \**p*<0.05, \*\**p*<0.01, \*\*\**p*<0.001, \*\*\*\**p*<0.001.

### Unilateral hippocampal injection reveals Aβ antibody within neurons and activation of astrocytes and microglia in APP*^NL-F^* mice

We next injected 6E10 into the hippocampus of pre-plaque, 5-month-old APP*^NL-F^* mice and examined the anatomical and cellular localization of Aβ antibody before plaque formation (Fig. 2a). Following unilateral IH injections, antibody labeling was detected in the injected side at both 24 h and 72 h post-injection (Fig. 2b, c). Compared with our results with the 5xFAD mice, above, which have abundant plaques, we found less labeling in the injected hippocampus of pre-plaque APP*^NL-F^* mice (Figure 2d). Remarkably, however, 24 h after Aβ antibody injection, we detected a clear signal inside of neurons, particularly in hippocampal field CA1 (Fig. 2e, h) and in a subset of neurons located in the deep layer of the CA1/subiculum transitional region (Fig. 2k). MOAB-2 labeling of the same sections for intracellular Aβ (Fig. 2f, i, l) revealed that the 6E10 antibody colocalized with intracellular Aβ in neurons (Fig. 2g, j, m). We also detected activated astrocytes (Fig. 2n, o, r) (uninjected-24 h vs injected-24 h, *p*=0.0001, uninjected-72 h vs injected-72 h, *p*<0.0001) and microglia (uninjected-24h vs injected-24 h, *p*<0.0001, uninjected-72 h vs injected-72 h, *p*=0.0472) (Fig. 2p, q, s) in these injected pre-plaque mice. The marked effect on astrocytosis with the very low p values likely, as noted for the 5xFAD mouse above, the fact that the needle injection itself induces robust astrocytosis. Of note, Aβ antibody 6E10-labeled microglia were widely distributed within the hippocampus (Fig. S1f, g). However, given the damage to the brain from the needle track, it is possible that the activation and antibody distribution in microglia are also not solely Aβ antibody-mediated. To avoid damage to brain parenchyma, we next, directly delivered Aβ antibody into the CSF via the cisterna magna (CM). In contrast to CM injection, the brain delivery efficiency of Aβ antibodies administered by intraperitoneal (IP) injection is relatively low. Previous studies have reported that only approximately 0.1–0.2% of circulating antibodies can cross the blood–brain barrier (BBB) and enter the brain[52]. Thus, to further investigate the effects of Aβ antibodies and to mitigate the potential effects of mechanical damage from the injection itself, we turned to antibody injection into the CM. Although CM injection of Aβ antibodies is a relatively invasive method, it allows for high concentrations of Aβ antibody in the CSF, thereby increasing brain parenchymal delivery and overall bioavailability.

**Fig. 2.**
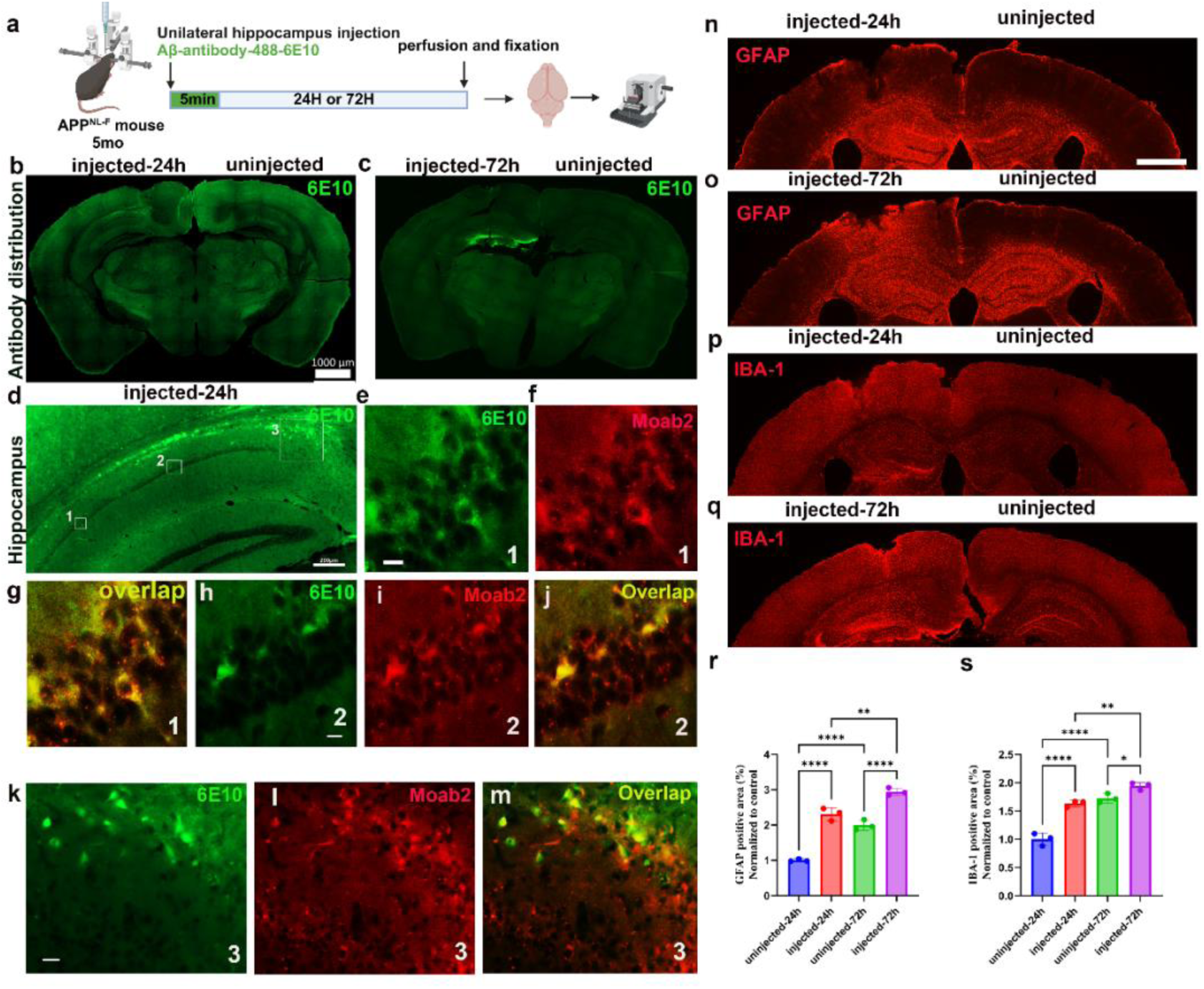
Unilateral intrahippocampal injection of Aβ antibody 6E10 in APP*^NL-F^* mice results in intraneuronal antibody in neurons and activation of astrocytes and microglia before the appearance of plaques. (a) 5 μl of Aβ antibody 6E10 was injected into the hippocampus of 5-month-old APP*^NL-F^* mice. (b, c) Representative images of Aβ antibody 6E10 at 24 and 72 h; scale bar = 1 mm. (d) Three areas were selected in CA1 and in a subset of neurons located in the deep layer of the CA1/subiculum transitional region for analysis; scale bar = 200 μm. (e, h, k) All selected areas showed intracellular localization of the Aβ antibody 6E10 in neurons; scale bar = 10 μm. (f, i, l) MOAB-2 labeling for intracellular Aβ of the same subregion as in e, h, and k; scale bar = 10 μm. (g, j, m) Injected Aβ antibody 6E10 colocalizes with MOAB-2 positive intracellular Aβ in the merged image; scale bar = 10 μm. GFAP levels in the hippocampus at 24 (n) and 72 h (o) after injection; scale bar = 1 mm. IBA-1 levels in the hippocampus 24 (p) and 72 h (q) after injection; scale bar = 1 mm. Quantification of the areas of the hippocampus covered by GFAP (r) and IBA-1(s) labeling; N=3 per group. Data are represented as mean ± SD. \**p*<0.05, \*\**p*<0.01, \*\*\**p*<0.001.

### Cisterna magna injection reveals Aβ antibody in different cell types of APP*^NL-G-F^*and APP*^NL-F^* mice

Unlike IH injection, CM injection should eliminate the potential for mechanical damage to the brain parenchyma, and Aβ antibody injected in CM will be able to distribute more broadly across the brain, though at lower local concentrations compared to direct intraparenchymal injection. Compounding this, a substantial portion of the CM-injected antibody will flow out of the brain as it travels with the circulating CSF. In order to enhance the translocation of Aβ antibody into the parenchyma, we tested an approach used in a previous study, wherein IP injection of hypertonic saline(HS) was shown to enhance the entry of CM-injected antibodies into the brain parenchyma[53]. To better evaluate the effect of IP injection of HS on CSF influx, we performed CM injection of bovine serum albumin conjugated to Alexa Fluor 647 (BSA-647) in 4-month-old wild-type mice (WT) and of Aβ antibody 6E10 in 4-month-old APP*^NL-G-F^* mice. Concurrently, we IP administered either isotonic saline (normal saline (NS), 0.154 M, IP-NS) or HS (1 M, IP-HS) (Fig. S3a, b). We found that in WT mice, the distribution in the brain parenchyma of BSA-647 in the IP-HS group was approximately 2.323 times greater than in the IP-NS group (*p*<0.0001). In APP*^NL-G-F^* mice, the distribution in the brain parenchyma of the 6E10 antibody in the IP-HS group was 2.485 times that of the IP-NS group (*p*=0.0003) (Fig. S3a, b, c, d, e, f). 3D-reconstruction revealed that IP-HS significantly enhanced antibody 6E10 localization around blood vessels and cortical plaques (Fig. S3g, h), suggesting improved antibody translocation with IP-HS. Henceforth, IP-HS injection accompanied all CM antibody injection experiments.

We then tested whether antibody 6E10 would have an effect in APP*^NL-G-F^* mice 6 h after injection into CM. To do this, mice were injected in CM with either 6E10 or PBS in addition to IP-HS injection (Fig. 3a). We found that 6E10 distributed widely throughout the brain (Fig. 3b), including in the hippocampus (Fig. 3c). Notably, like with IH antibody injection, CM injection resulted in antibody accumulation within neurons of CA1 (Fig. 3d) and subiculum (Fig. 3g), wherein it colocalized with intracellular Aβ shown by MOAB2 antibody (Fig. 3e, f, h). CM injection of 6E10 also significantly reduced cortical plaques (cortex: *p*=0.0418, hippocampus: *p*=0.7799) (Fig. S4a, b, e), and activated microglia (cortex: *p*=0.0168, hippocampus: *p*=0.0017) (Fig. S4c, f) and astrocytes (cortex: *p*=0.0004, hippocampus: *p*=0.6172) (Fig. S4d, g) when combined with IP-HS. However, unlike at 72 h post-IH injection in 5xFAD mice, in APP*^NL-G-F^* mice, there was no significant reduction in intracellular Aβ in CA1 neurons of the hippocampus at this shorter 6 h post injection time with 6E10 compared to PBS (Fig. S2g, h, j, k, l).

**Fig. 3.**
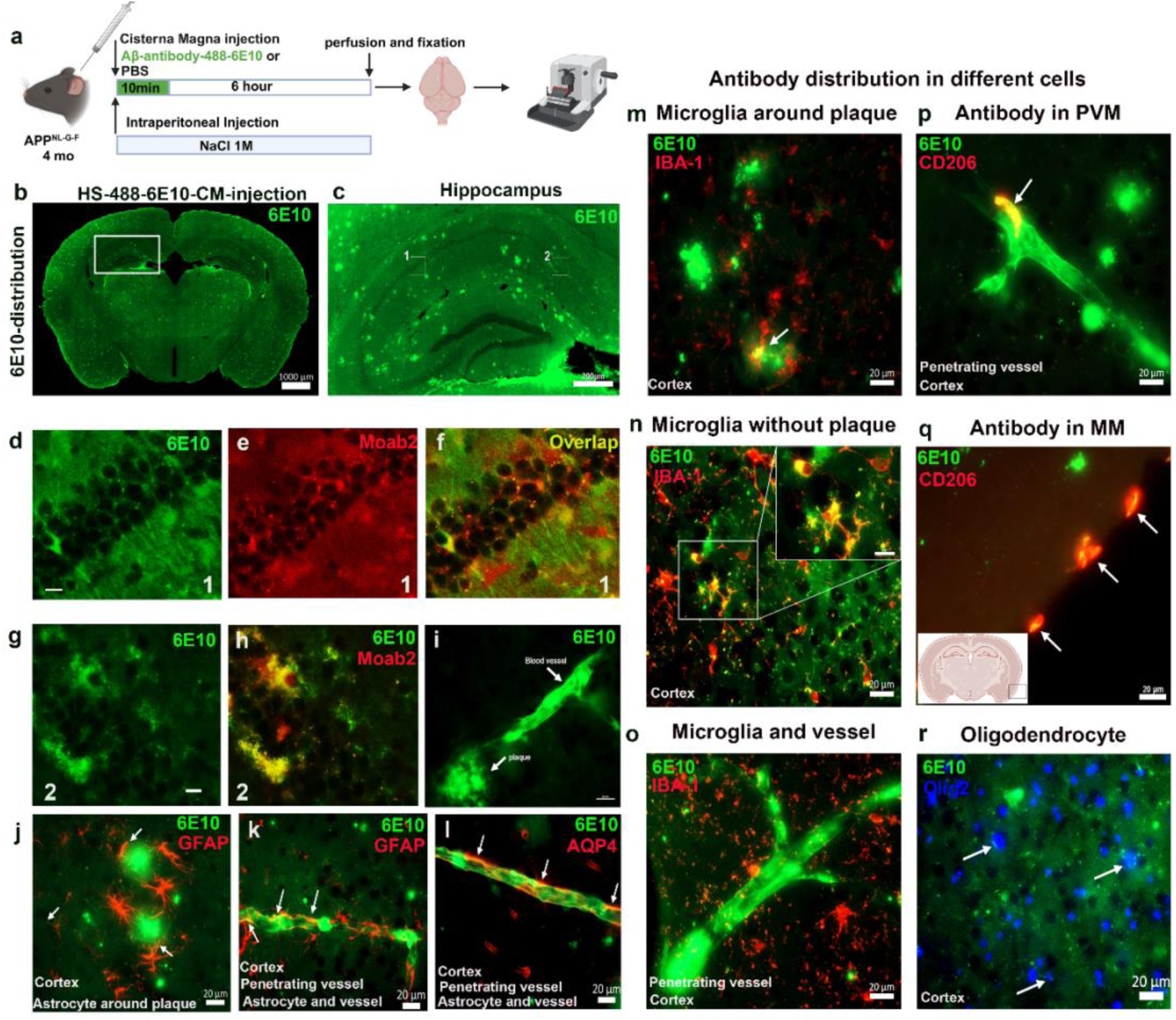
Aβ antibody 6E10 is detected in different brain cells following CM injection in 4-month-old APP*^NL-G-F^* mice. (a) 10 μl of Aβ antibody 6E10 or PBS was injected into the CM combined with an IP injection of hypertonic saline. (b) The distributions of Aβ antibody 6E10 in the brain; scale bar = 1 mm. (c) The distributions of Aβ antibody 6E10 in the hippocampus; scale bar = 200 μm. (d) Aβ antibody 6E10 is detected in CA1 neurons; scale bar = 10 μm. (e, f) Antibody MOAB-2 labeling in CA1 shows that intraneuronal Aβ colocalizes with the injected Aβ antibody 6E10; scale bar = 10 μm. (g) The Aβ antibody 6E10 in subiculum neurons; scale bar = 10 μm. (h) Antibody MOAB-2 labeling in subiculum shows that intraneuronal Aβ colocalizes with Aβ antibody 6E10. (i) Aβ antibody 6E10 in penetrating vessels and plaques. Aβ antibody 6E10 in astrocytes around plaques (j), and in penetrating vessels (k) in the cortex; scale bar = 20 μm. (l) Aquaporin-4 (AQP4) labeling reveals Aβ antibody 6E10 in astrocytic endfeet around blood vessels in the cortex; scale bar = 20 μm. Aβ antibody 6E10 in microglia around plaques (m) and in areas without plaques (n) in the cortex; scale bar = 20 μm. Aβ antibody 6E10 accumulates in the microglia around blood vessels (o), perivascular macrophages (PVMs) (p) of penetrating vessels, and meningeal macrophages (MM) (q), and oligodendrocytes (r); scale bar = 20 μm. N=4 per group.

To further investigate where else antibody 6E10 localized, we examined glial cells near penetrating cortical vessels and large arteries. Consistent with the unilateral IH injections, we detected a substantial amount of antibody surrounding blood vessels and associated with plaques (Fig. 3i). Astrocytes adjacent to plaques exhibited an activated morphology (astrocytes undergo morphological hypertrophy, characterized by enlarged cell bodies and thickened processes compared to the control group) but showed only minor colocalization with the antibody (Fig. 3j). However, we noticed that in cortical penetrating vessels, astrocytes closely associated with the vasculature contained a relatively greater amount of antibody than the astrocytes around plaques (Fig. 3k). To further examine this, we utilized an antibody against aquaporin-4 (AQP4), a marker of astrocytic end feet, and observed colocalization between AQP4 and the injected Aβ antibody around blood vessels (Fig. 3l). Next, we analyzed the colocalization of antibody 6E10 with microglia in different cortical regions, including plaque-burdened (Fig. 3m) and plaque-free cortical areas (Fig. 3n). We observed a substantial amount of 6E10 inside microglia. Surprisingly, this appeared to be independent of the presence of plaques in these AD mice.

Aβ is known to accumulate in blood vessels and cause cerebral amyloid angiopathy (CAA) in AD[54] and a main side effect of Aβ immunotherapies for AD is microhemorrhage[49]. Because of this, we further investigated Aβ antibody localization around blood vessels. Interestingly, perivascular microglia (Fig. 3o) and perivascular macrophages (PVMs) showed accumulation of Aβ antibody 6E10 (Fig. 3p, Fig. S5a, b, c). Additionally, we detected 6E10 signal in meningeal macrophages (Fig. 3q) and in oligodendrocytes (Fig. 3r); in the latter we detected the injected Aβ antibody within the cytoplasm surrounding nuclei labeled with oligodendrocyte-specific antibody Olig2. Oligodendrocytes were also found to contain Aβ42, which localized in the cells with the intracellular Aβ antibody 6E10 (Fig. S5d, e, f, g). Together, our results show a remarkably widespread distribution of injected antibody 6E10 across various cell types within the brain.

We next examined the distribution of CM-injected Aβ antibody 6E10 in the mouse brain before the appearance of plaques (Fig. S6a). Pre-plaque 5-month-old APP*^NL-F^*mice showed that 6E10 was widely distributed throughout the brain (Fig. S6b), including the hippocampus (Fig. S6c). Some 6E10 antibody was found in neurons of CA1 (Fig. S6d) and CA3 (Fig. S6g), wherein a portion co-localized with intracellular Aβ labelled with MOAB-2 (Fig. S6e, f, h, i). Furthermore, we observed injected 6E10 in PVMs (Fig. S6j), oligodendrocytes (Fig. S6k), and microglia (Fig. S6l).

In comparison, the distribution of CM-injected 6E10 in 5-month-old WT mice differed from that in AD mouse models (Fig. S6m, n). We noted only a small fraction of 6E10 was associated with blood vessels, and there was no detectable injected Aβ antibody in neurons of the hippocampus. However, we did observe 6E10 in PVMs (Fig. S6o) and microglia (Fig. S6p) of WT mice, which we speculate may be related to the phagocytic function of these two cell types.

### Aβ antibody 6E10 distribution in blood vessels following CM injection

As indicated above, Aβ antibody 6E10 injected in CM accumulated heavily around blood vessels in APP*^NL-G-F^* mouse brain. To determine more precisely where in the vessels the injected antibody 6E10 was localized, we labeled with markers against different parts of the vessel structure, namely CD31, elastin and α-SMA. Firstly, labeling with CD31, which is primarily located on the surface of vascular endothelial cells, showed no colocalization with Aβ antibody 6E10 (Fig. 4a). To further investigate antibody 6E10 localization within blood vessels, we used a stain for elastin, a marker for the vascular basement membrane. The endothelial cell CD31 antibody localized inside of elastin and 6E10 antibody outside of elastin (Fig. 4b). Antibody 6E10 is also outside of the vascular smooth muscle layer labeled using an antibody to α-SMA (Fig. 4c). With the α-SMA labeling, we were able to detect the presence of 6E10 also in large vessels of the circle of Willis (Fig. 4d). There, α-SMA-positive smooth muscle cells were observed surrounding the CD31-positive endothelial cells at a distance, with laminations of 6E10, though the injected Aβ antibody primarily accumulated in the outermost layer of these vessels. Lastly, we labeled with a C-terminus-specific Aβ40 antibody, which is associated with blood vessels in CAA in AD. We found that Aβ40 mainly localized outside the vascular endothelium of vessels of the circle of Willis, although most of the injected 6E10 signal did not colocalize with the Aβ40 labelling (Fig. 4e). However, aggregated forms of Aβ are not effectively detected by C-terminus-specific Aβ antibodies [55].

**Fig. 4.**
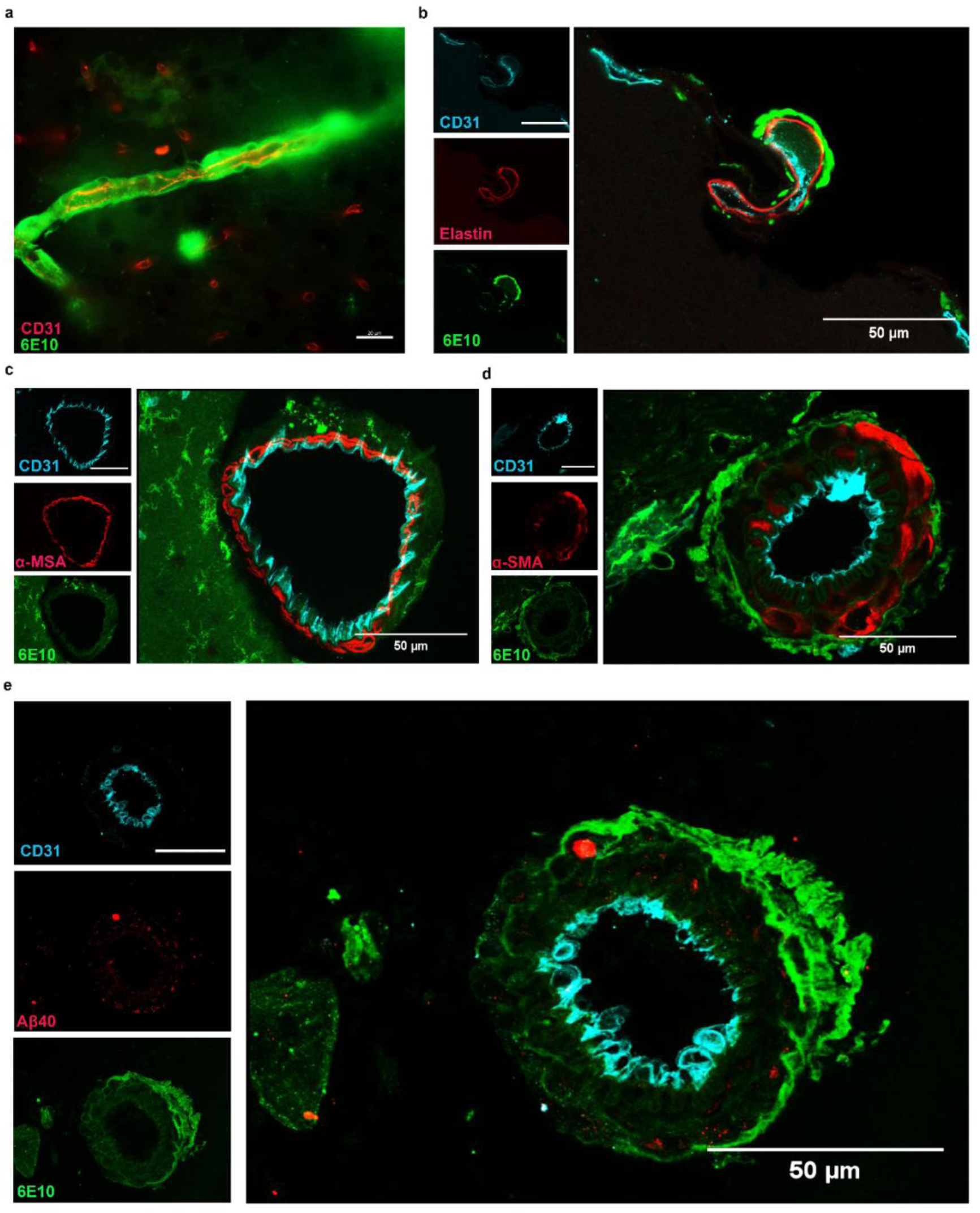
The distributions of Aβ antibody 6E10 in penetrating vessels and large arteries of 4-month-old APP*^NL-G-F^* mice following CM antibody injection. (a) Labeling with the endothelial cell marker CD31 shows that the Aβ antibody 6E10 is primarily located in the outer layer of the vascular endothelium; scale bar = 20 μm. (b) Aβ antibody 6E10 is distributed outside of the elastin layer; scale bar = 50 μm. (c) Aβ antibody 6E10 is distributed on the outer side of the vascular smooth muscle layer; scale bar = 50 μm. (d) In large arteries of the circle of Willis, vascular smooth muscle cells are located on the outer layer of the endothelium but at a greater distance, while Aβ antibody 6E10 is distributed in the outermost layer of the blood vessels; scale bar = 50 μm. (e) In large vessels of the circle of Willis, the localization of Aβ40 is different from that of injected Aβ antibody 6E10 within blood vessels; scale bar = 50 μm. N=4 per group.

### IP injection reveals Aβ antibody in brain cell types and blood vessels of APP*^NL-G-F^* mice

In clinical settings, Aβ antibodies such as lecanemab or donanemab are administered intravenously to AD patients, whereas in preclinical AD mouse models, they are typically delivered via IP injection[1]. We used IP injection of 6E10 (2 mg/kg) to investigate its distribution in different brain cell populations and blood vessels. A single IP injection of 647–6E10 was administered, and tissue samples were harvested 3 days post-injection (Fig. 5a). We found that antibodies administered via IP injection were widely distributed in the brain (Fig. 5b). Remarkablsy and unlike CM injection, after IP injection antibodies were not only detected in penetrating vessels but also in capillaries of the brain parenchyma. High-magnification views of the hippocampus revealed clear Aβ antibody binding to amyloid plaques as well as distribution within capillaries in the hippocampus (Fig. 5c). Moreover, we observed intracellular accumulation of Aβ antibody particularly in neurons located in the CA1 region. We selected two regions for high-magnification analysis to more clearly show antibody 6E10 localized within neurons in the CA1 regions (Fig. 5d, g) and co-localizing with Aβ42 (Fig. 5e, f, h, i). Some IP-injected Aβ antibody was also detected in astrocytes surrounding plaques (Fig. 5j). However, unlike with CM injection in perivascular astrocytes, the IP-injected Aβ antibody was predominantly confined within the blood vessels and did not co-localize with astrocyte endfeet (Fig. 5k). Aβ antibody was localized in microglia both near and distant from plaques (Fig. 5i) and surrounding the nuclei (olig2) of oligodendrocytes (Fig. 5m). We next examined cortical penetrating vessels (Fig. 5n) and small hippocampal vessels (Fig. 5o) to investigate Aβ antibody distribution in PVMs, and observed a prominent accumulation of the Aβ antibody within these cells. To more precisely determine the vascular localization of IP-injected antibody6E10, we examined cortical penetrating vessels (Fig. 5p, r), small hippocampal vessels (Fig. 5q), and arteries in the circle of Willis (Fig. 5s). The Aβ antibody localized to the abluminal perivascular compartment, lying outside the endothelial (CD31) and vascular smooth muscle (α-SMA) layers but inside the boundary formed by astrocytic endfeet. Although Aβ antibodies administered via IP injection cross the blood–brain barrier (BBB) to reach the brain parenchyma, Aβ antibody was not detected in the vascular endothelium or vascular smooth muscle layer.

**Fig. 5.**
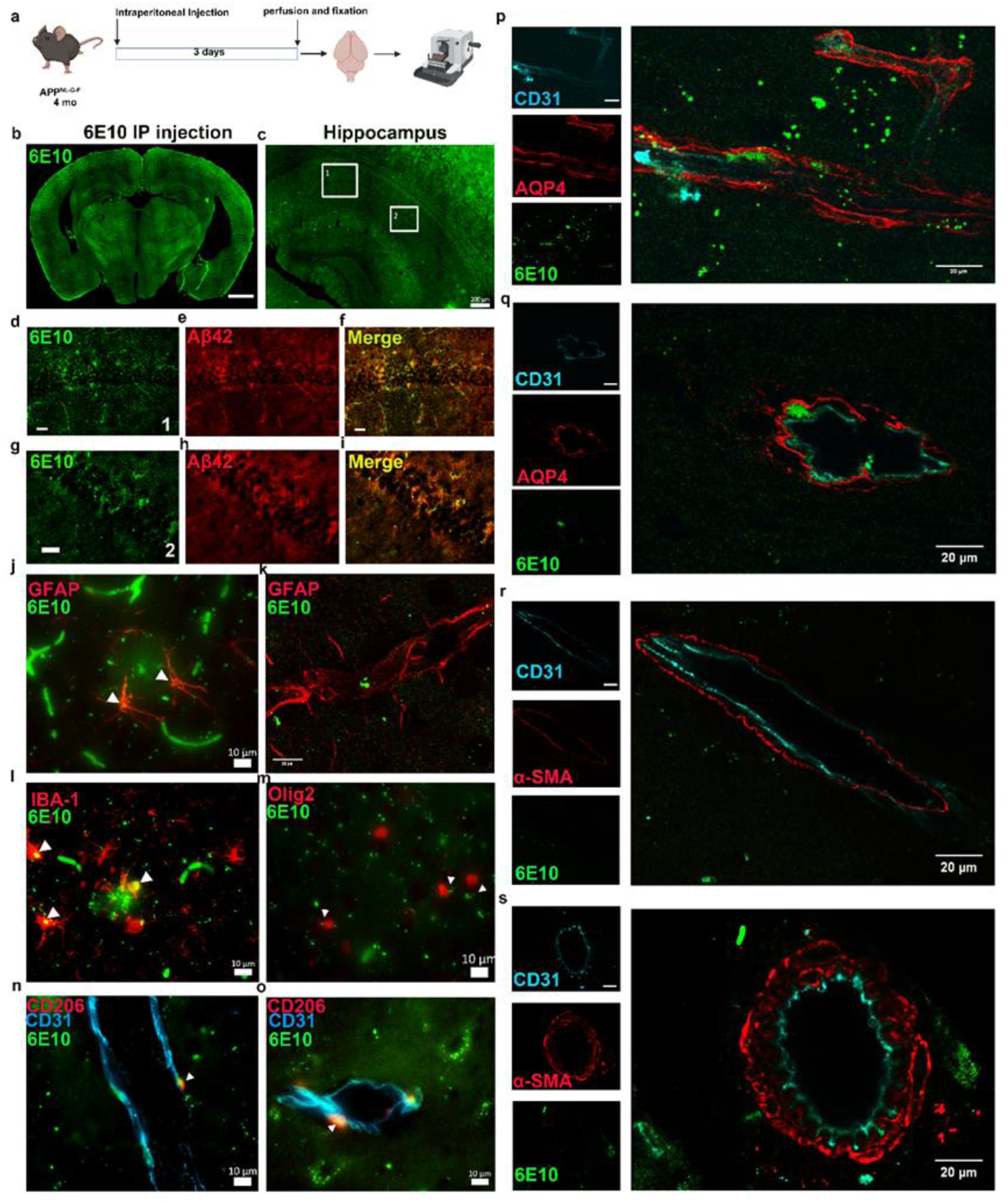
Aβ antibody 6E10 is detected in different brain cells following IP injection in 4-month-old APP*^NL-G-F^* mice. (a) Mice received a single IP injection of Aβ antibody 6E10 at a dose of 2 mg/kg, and tissues were collected 3 days post-injection. (b) The distributions of Aβ antibody 6E10 in the brain; scale bar = 1 mm. (c) The distributions of Aβ antibody 6E10 in the hippocampus, two areas were selected in CA1 for analysis; scale bar = 200 μm. (d, g) Aβ antibody 6E10 is detected in CA1 neurons; scale bar = 20 μm. (e, f, h, i) Antibody Aβ42 labeling in CA1 shows that intraneuronal Aβ colocalizes with the injected Aβ antibody 6E10; scale bar = 20 μm. (j) Aβ antibody 6E10 in astrocytes around plaques, but not localized to the pericascular astrocytes (k). Aβ antibody 6E10 in microglia near and distant from plaques in the cortex (i). Aβ antibody 6E10 surrounds the nuclei of oligodendrocytes (m). Aβ antibody 6E10 in PVMs in a penetrating vessel (n) in the cortex and small vessel (o) in the hippocampus; scale bar = 10 μm. The Aβ antibody 6E10 is distributed outside the endothelial cell layer (CD31) but inside astrocytic endfeet (AQP4) in the penetrating vessel (p) and small vessel in the hippocampus (q); scale bar = 20 μm. The Aβ antibody 6E10 is distributed outside CD31 and smooth muscle cell layer (α-SMA) in the penetrating vessel (r) and vessels originating from the circle of Willis (s); scale bar = 20 μm. N=2 per group.

### Effect of Aβ antibody 6E10 on the glymphatic system

Finally, we examined whether injection of 6E10 would affect glymphatic circulation. To do this, we performed two subsequent injections in 5xFAD mouse brain: a unilateral IH injection of Aβ antibody 6E10 and a CM injection with BSA-647, but without IP-HS, either 24 h or 72 h post-IH antibody injection (Fig. 6a). We did not use IP-HS for these experiments, since we wanted to evaluate effects of antibody on BSA influx, which would be strongly affected by IP-HS. No significant changes in BSA-647 influx were seen within the antibody injected hippocampus when compared to the uninjected side at both the 24 h and 72 h timepoints (uninjected-24 h vs injected-24 h, *p*=0.2135, uninjected-72 h vs injected-72 h, *p*=0.8832) (Fig. 6b, c, Fig. S7a for analysis). To extend the circulation time of the Aβ antibody in the brain, we next performed unilateral intracerebroventricular (ICV) injection of 6E10 in APP*^NL-G-F^*mice and allowed it to circulate for 24 h before CM injection of BSA-647 without IP-HS (Fig. 6d). ICV Aβ antibody injection in APP*^NL-G-F^* mouse brains also did not significantly alter glymphatic system circulation compared to the ICV PBS group (*p*=0.2) (Fig. 6e, f, Fig. S7b for analysis).

**Fig. 6.**
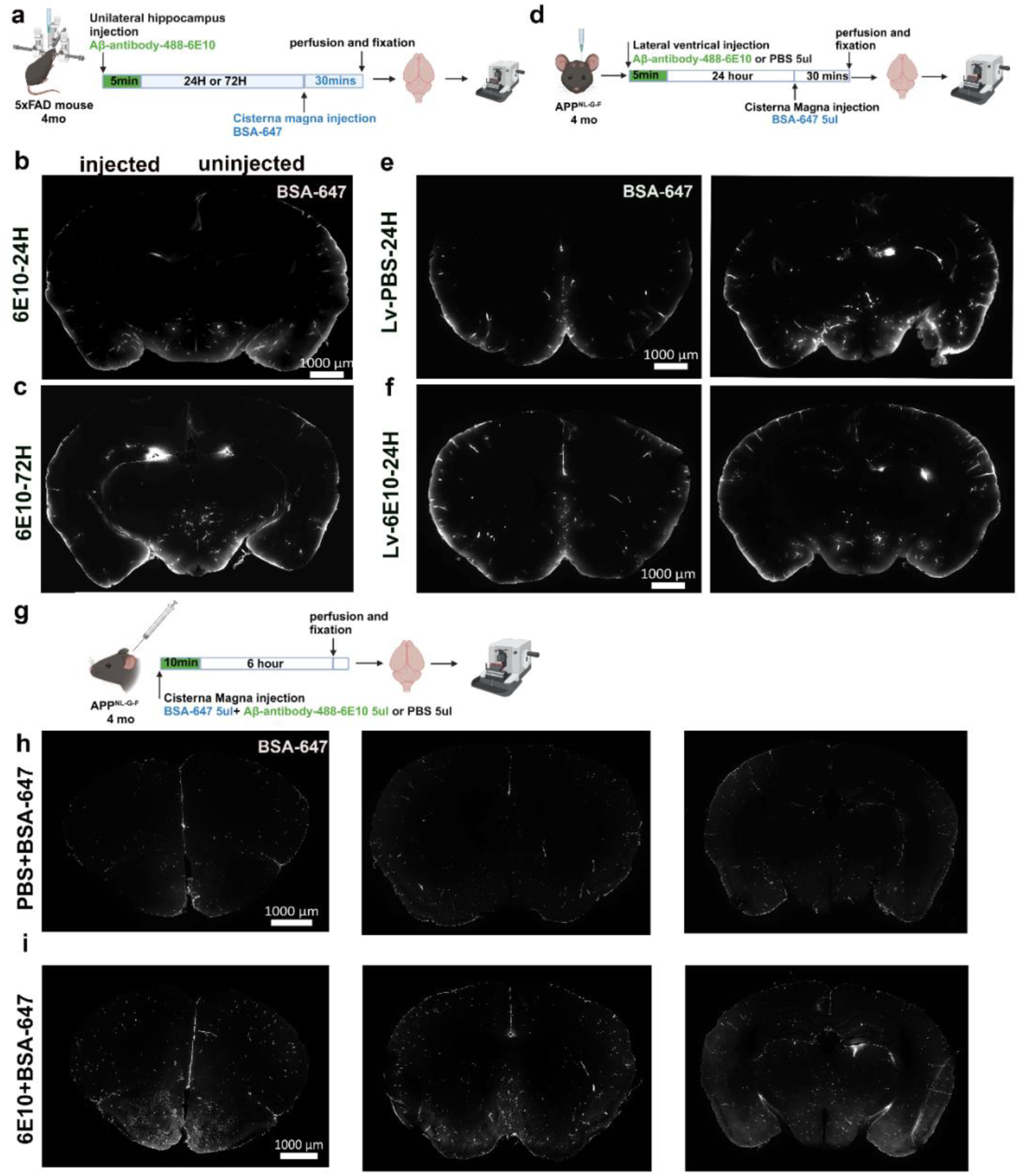
Effects of Aβ antibody 6E10 injection on glymphatic function. (a) Unilateral IH injection of 5 μL of Aβ antibody 6E10 in 4-month-old 5xFAD mice, followed by CM injection of 5 μL of BSA-647. (b, c) Representative images of BSA-647 distribution after unilateral IH injection of Aβ antibody 6E10 at 24 and 72 h; scale bar=1 mm. (d) A total of 5 μL of Aβ antibody 6E10 or PBS was injected into the lateral ventricle of 4-month-old APP*^NL-G-F^* mice, followed by injection of 5 μL of BSA-647 in CM. (e, f) Representative images of BSA-647 distributions after lateral ventricle injection of Aβ antibody 6E10 or PBS at 24 h; scale bar = 1 mm. (g) Co-injection of 5 µL BSA-647 and 5 µL Aβ antibody 6E10 or PBS in the CM, and the BSA-647 distribution in the brain was examined after 6 h. (h, i) Representative images of BSA-647 distributions after co-injection of Aβ antibody 6E10 or PBS with BSA-647; scale bar = 1 mm.

To control for potential mechanical damage caused by IH or ICV injections, we next injected both 6E10 and BSA-647 exclusively into CM in another group of APP*^NL-G-F^* mice. Typically, glymphatic system function is assessed by allowing BSA-647 to circulate for 30 minutes[53], but to maintain consistency with our previous Aβ antibody circulation experiments, we allowed the BSA-647 and 6E10 mixture to circulate in the brain for 6 h before examining BSA-647 distribution (Fig. 6g). We found that with co-injection into CM, Aβ antibody 6E10 did not affect glymphatic circulation in 4-month-old APP*^NL-G-F^* mice, although there was a trend towards an increase in the accumulation of vessel-associated BSA-647 in brain sections (*p*=0.1753) (Fig. 6h, i, Fig. S7c for analysis). For these co-injection experiments of Aβ antibody 6E10 and BSA-647, we separately show the distribution of BSA-647 (Fig. S7d), Aβ antibody 6E10 (Fig. S7e), and PVMs (Fig. S7f) in the brain. To analyze where in the brain this BSA-647 was, we used CD206 antibody for PVMs and detected that the accumulated BSA-647 colocalized with CD206 antibodies for PVMs (Fig. S7g, h, i). CD206 antibody labelling also revealed macrophages harboring BSA-647 in cortical penetrating vessels and small blood vessels of the hippocampus (Fig. S7g, h, i).

## Discussion

In this study we aimed to investigate the more precise distribution of the N-terminal Aβ domain antibody 6E10 in brain, following its injection via different routes, including its potential association with different brain cells and blood vessels. We additionally examined potential effects on the glymphatic system. We show that this widely used Aβ antibody in the AD research field localizes not only to plaques and blood vessels, but also to different cell types in the brain, including neurons, astrocytes, microglia, perivascular macrophages, and oligodendrocytes. In hippocampal field CA1 and the subiculum, including in their border region, Aβ antibody 6E10 localized with intracellular Aβ in these AD-vulnerable neurons known to be prone to early death in AD and to contain more intraneuronal Aβ[56]. In astrocytes with CM but not IP injections, antibody 6E10 predominantly localized to their endfeet abutting blood vessels, while in microglia, antibodies localized to the cell soma. Further, with respect to cerebral blood vessels, we observed that Aβ antibody 6E10 primarily localized to the outermost layer, but that it did not colocalize much with antibodies against the free C-terminus of Aβ40. However, a caveat with this observation is the known propensity for aggregating Aβ to undergo conformational changes that typically obscure labeling with C-terminus Aβ antibodies, meaning that this labeling should be interpreted with caution. Finally, our results indicate that Aβ antibody 6E10 did not appear to alter the glymphatic circulation.

Previous studies have demonstrated that Aβ antibodies presented to the brain can reduce plaques, activate astrocytes and microglia, and enhance the phagocytic activity of microglia[57–59]. Elevated levels of intracellular Aβ in neurons are increasingly viewed as an important contributor to cognitive impairment and synaptic damage in AD mouse models [11, 60, 61], and its effects are also being investigated by way of mathematical modelling[62]. Active immunization with the Aβ derivative K6Aβ1–30-NH₂ in a non-human primate model resulted in a marked reduction of intraneuronal Aβ accumulation in the brains of mouse lemurs[63]. Our prior mechanistic cellular work had shown that Aβ antibodies could be internalized into neurons in culture but not that this can also occur in vivo[27]. In ongoing studies we are continuing to examine the mechansisms of antibody internalization and cellular Aβ reduction, and also how approved Aβ therapeutic antibodies compare to the results obtained with the N-terminal Aβ antibody employed in this study.

A recent study showed that the deletion of Beta-site APP Cleaving Enzyme 1 (BACE1) in oligodendrocytes reduces Aβ plaque burden in an AD mouse model[64]; such studies have suggested that oligodendrocytes might also be active contributors to Aβ pathology in AD[64, 65]. Our results show that oligodendrocytes also contain injected Aβ antibody that localizes with intracellular Aβ, which may provide a rationale for considering oligodendrocytes as potential therapeutic targets. Due to their potent phagocytic capacity, PVMs are closely associated with the clearance of systemically administered Aβ antibodies around blood vessels. Further investigation is needed to explore potential effects on PVMs after uptake of Aβ antibody. We found a rapid micro- and astro-glial response already 6 h after administering Aβ antibodies (using hypertonic IP injections). In view of the importance of neuroinflammation in AD, further studies are warranted to explore this glial antibody response and also its potential effect on neurons. In our study, we found that the Aβ antibody did not alter glymphatic function in AD mice. This may however be affected by structural disruption of the brain caused by the injection methods used.

ARIA is the most important side effect of both currently FDA approved amyloid immunotherapies Lecanemab and Donanemab, but the molecular mechanism(s) and which cell types are most involved remain to be fully understood. We show that Aβ antibody localizes to the outermost layer of blood vessels as well as within perivascular macrophages, astrocytic endfeet, and microglia. Further studies are needed to explore the effects of Aβ antibodies on these cell types in relation to vascular amyloid.

## Conclusions

Antibodies are increasingly used to target also intracellular proteins, including, for example, the microtubile-associated protein tau for AD therapy[66]. Our study underscores that a widely used Aβ antibody exhibits a broad distribution profile in various cell types within the brains of different AD mouse models. Beyond its localization to amyloid plaques, antibodies were detected in neurons, microglia, astrocytic endfeet, perivascular macrophages, and oligodendrocytes. These findings indicate that, in the context of immunotherapy, antibody distribution is not limited to amyloid plaques and may influence various cell populations, which could represent novel therapeutic targets in future immunotherapeutic strategies for AD.

## Materials and Methods

### Animals

Male mice were allowed a minimum acclimation period of five days before any handling or experimental procedures. They were housed in a temperature- and humidity-controlled environment with a 12-hour light/dark cycle, in groups of three to five per individually ventilated cage. All cages were enriched with toys to promote animal welfare, and animals had ad libitum access to standard chow and water. All mouse experiments were ethically approved by the Malmo/Lund Ethics Committee on Animal Testing (dnr 5.8.18-13038/2024). For antibody injections, we used several different AD mouse models. 1. 5xFAD mice (Jackson Laboratory, B6SJL-Tg [APPSwFILon, PSEN1*M146L*L286V]6799Vas/Mmjax), The familial Alzheimer’s disease (FAD) mutations APP K670N/M671L (based on the 770-residue isoform), along with I716V and V717I, as well as PS1 mutations M146L and L286V, were introduced into the cDNAs of APP (695) and PS1 through site-directed mutagenesis; these modified sequences were then subcloned into exon 2 of the mouse Thy1 transgene cassette, as described[14], and subsequently verified by sequencing using standard protocols. 2. APP *^NL-G-F^* mice (Jackson Laboratory, APPtm3.1Tcs) harbor mutations: APP K670_M671delinsNL (Swedish), APP I716F (Iberian), and APP E693G (Arctic); these mutations promote Aβ pathology by increasing total Aβ production (Swedish mutation), increasing the Aβ_42_/Aβ_40_ ratio (Iberian mutation), and promoting Aβ aggregation through facilitating oligomerization and reducing proteolytic degradation (Arctic mutation)[67]. 3. APP*^NL-F^*mice (Jackson Laboratory, APPtm2.1Tcs); for the APP*^NL-F^* knock-in mice, the APP construct contains a humanized Aβ region along with two pathogenic mutations, the Swedish “NL” and the Iberian “F”[67]. Due to the use of the endogenous mouse APP promoter, the constructs of the APP*^NL-F^* and APP *^NL-G-F^* mice are expressed in appropriate cell types and temporal specificity.

### Experimental time points

Antibody 6E10 is an IgG1 mouse monoclonal antibody that has been widely used in AD research and is directed at the N-terminal domain of Aβ, and sees Aβ peptides and Aβ-containing full-length amyloid precursor protein (APP) and βAPP. C-terminal fragment. Experimental research has shown that Aβ N-terminal domain antibodies reduce amyloid plaques and improve behavior in AD mice[68, 69]. For 5xFAD mice and APP*^NL-F^* mice, unilateral hippocampal injection of Alexa Fluor 488 conjugated 6E10 antibody was performed at 24 and 72 h before sacrifice. We used the guinea pig anti-rabbit Alexa Fluor 647 conjugated secondary antibody for a control group for the 5xFAD mouse intrahippocampal injections. Mice were anesthetized with isoflurane and then underwent perfusion with phosphate buffered saline (PBS) 0.1 M, pH 7.4, followed by fixation with 4% freshly depolymerized paraformaldehyde (PFA). Subsequently, their brains were immersed in 4% PFA overnight. Additionally, another group of 5xFAD mice receiving intrahippocampal 6E10 antibody injection 24 h and 72 h before being sacrificed underwent cisterna magna (CM) injections of BSA-647. After 30 minutes of BSA-647 circulation, these latter mice were anesthetized with IP injection of a mixture containing ketamine (100 mg/ml) and domitor (1 mg/ml) at a dose of 10 μl/g, followed by perfusion with 0.1M PBS and 4% PFA. To assess the effect of hypertonic saline, WT or APP*^NL-G-F^* mice were administered an IP injection of either normal (0.154 M) or hypertonic (1 M) saline concurrently with a CM injection of 10 μl of BSA-647 or 6E10 antibody. BSA-647 or 6E10 antibodies were freely circulating for 30 mins after injection into the CM. Mice were then perfused with PBS and 4% PFA. We found that hypertonic saline IP injections allowed for more 6E10 antibodies to be localized in the brain. Thus, we used hypertonic saline IP injection combined with the 6E10 antibody CM injection in the later experiments and allowed the 6E10 antibody to freely circulate for 6 h. For the lateral ventricle injections with 6E10 antibody in APP*^NL-G-F^* mice, CM injections of BSA-647 were performed after 24 h, with BSA-647 circulation for 30 mins, and then the mice were sacrificed to evaluate glymphatic influx. The mixture of 6E10 plus BSA-647 CM injected into APP*^NL-G-F^*mice was left to circulate for 6 h before the mice were perfused following the method above. For the 6E10 antibody conjugated with Alexa Fluor 647 IP injection, APP*^NL-G-F^* mice were sacrificed, and tissues were collected 3 days post-IP injection. Brains were carefully excised from the skull and placed in PFA at 4 ℃ overnight. Then, brain sections were prepared using a microtome and stored in cryoprotective solution (30% sucrose and 30% ethylene glycol in 0.1M PB).

### Intracerebral injection

#### Intrahippocampal injection

Unilateral intrahippocampal (IH) injections were conducted following our previously published protocol[70]. In brief, mice were anesthetized using isoflurane (4% for induction, 2% for maintenance), and their heads were shaved and sterilized with 70% ethanol before being secured in a stereotactic apparatus. A surgical incision was made to expose the skull, and a small hole was drilled to accommodate the injection. The coordinates for the injections into the hippocampus were as follows: Bregma-2.5 mm, ML 2.0 mm, DV-1.8 mm. For antibody injections, 5 μl of 6E10 (purified anti-β-amyloid, Biolegend, #803003), or Alexa Fluor 488 anti-β-amyloid, 6E10 (clone 6E10, Biolegend, #803013), was injected at a speed of 1 μl/min via a Hamilton syringe (Hamilton, 7634-01). The needle was kept in place for 5 minutes to allow for diffusion of the antibody. The incision was then sutured, and the mice were monitored until they were awake from anesthesia.

#### Intracerebroventricular injection

The intracerebroventricular (ICV) injection protocol we used was as described in following refernece[71]. In brief, 6E10 antibody was injected directly into the right ventricle of the brain by using a stereotaxic apparatus. Before the surgery, their heads were shaved and sterilized with 70% ethanol before being secured in the stereotactic frame. The mice were anesthetized with isoflurane (4% for induction, 2% for maintenance). A surgical incision was made to expose the skull, and a small hole was drilled to accommodate the injection. The coordinates for the injections into the lateral ventricle was as follows: Bregma -0.2 mm, ML 1.0 mm, DV -2.3 mm below the brain surface. For injections, 5 μl of Alexa Fluor® 488 anti-β-Amyloid (clone 6E10, Biolegend, # 803013) were injected at a speed of 1 μl/min via a Hamilton syringe (Hamilton, 7634-01). The needle remained in position for 5 minutes to facilitate the diffusion of the antibody. Afterward, the incision was sutured, and the mice were observed until they had fully recovered from anesthesia.

### Cisterna magna injections

#### Antibody injections

Cisterna magna (CM) injections were conducted as previously reported[53]. In brief, mice were weighed and anesthetized with a mixture of ketamine (100 mg/ml) and domitor (1 mg/ml) at a dose of 10 μl/g by IP injection. Throughout the procedure, we ensured that the mice remained under deep anesthesia. The mice were secured in a stereotaxic frame, and the CM was surgically exposed. A 30G needle connected to a 25-µl Hamilton syringe (Hamilton, 80223) via a polyethylene tube was then inserted into the CM. The needle was secured in place using glue, and the solution containing the antibody was infused using a micro-injection pump (Science Products model nano jet, serial # 47927) at a rate of 1 μl/min. To safeguard against backflow of CSF, the needle was kept in place for 6 hours following the completion of the infusion. We placed the mice in an incubator to maintain their body temperature after the antibody injection.

#### BSA-647 injections

A solution of BSA-647 (bovine serum albumin, Alexa Fluor™ 647 conjugate, 66kDa, Invitrogen, A34785) was prepared by dissolving BSA-647 in artificial CSF at a ratio of 1:100. Using a micro-injection pump, this solution was injected into the CM at 1 μl/min under four conditions. In the first condition, the impact of hypertonic saline or normal saline on the BSA-647 distribution in the brain of WT mice was examined. We thus injected a 10 μl volume of the BSA-647 solution and allowed it to circulate for 30 minutes. In the second condition, using 5xFAD mice, 5 μl of the BSA-647 solution was injected into the CM, following either 24 h or 72 h after a unilateral IH injection of 6E10 antibody solution. The BSA-647 solution was again allowed to circulate for 30 minutes. In the third condition, using APP*^NL-G-F^* mice, we first injected 6E10 antibody solution into the lateral ventricle and then waited for 24 h, after which we injected 5 μl of the BSA-647 solution into the CM and let it circulate for 30 minutes. In the fourth condition, we injected the mix of 5μl 6E10 plus 5μl BSA-647 in the CM of APP*^NL-G-F^* mice and allowed the mix to circulate for 6 hours.

#### IP injection

4 months old APP *^NL-G-F^* mice were administered Alexa Fluor 647-conjugated 6E10 antibody (647–6E10) via IP injection at a dose of 2 µg/kg body weight. Tissue samples were collected 3 days post-injection.

#### Solutions

Solutions were prepared as previously reported[29, 53]. Normal saline (isosmotic:0.154 M NaCl in ddH_2_O; 20 μl/g). Hyperosmolality was induced with hypertonic saline (HS:1 M NaCl in ddH_2_O; 20 μl/g). Artificial CSF (aCSF) was composed of NaCl 124 mM, KCl 3 mM, NaHCO_3_ 26 mM, NaH_2_PO_4_ 1.24 mM, MgSO_4_ 2 mM, CaCl_2_ 2 mM, and Glucose 10 Mm. PB solution: 14.42g Na_2_HPO_4_*2H_2_O, 2.62g NaH_2_PO_4_*H_2_O in 1 L ddH_2_O.

#### Immunofluorescence

After fixation with 4% PFA, mouse brains were first immersed in a 15% and then in a 30% sucrose solution, each for 1 day. Subsequently, using a microtome, brains were cut into 40 μm coronal sections and stored in a cryoprotectant (30% sucrose and 30% ethylene glycol in 0.1 M PB, ph 7.4) until use. Sections were labeled using a free-floating method. First, sections were washed for 5 min × 3 in 0.1 M PBS. For labeling against Aβ, sections were treated with 88% formic acid in PBS for 8 mins and then washed 3 times for 5 min each with PBST (0.3% Triton X-100). Sections were then blocked for 1.5 h (3% normal donkey serum (NDS) or 3% normal goat serum (NGS) in PBST) at room temperature (RT), and then incubated in primary antibody containing the same blocking solution overnight at 4 ℃ on an agitator. After 3×10 min washes in PBS at RT, sections were then incubated with appropriate secondary antibodies for 2 h at RT and washed 3 times for 10 min each. Then, the sections were incubated for 5 min with 1:1000 DAPI. The antibodies used in this study were: Alexa Fluor 488 anti-β-amyloid, 6E10 (Biolegend, 803013, RRID: AB_2564765), Alexa Fluor 647 anti-β-amyloid, 6E10 (Biolegend, 803021, RRID: AB_2783374). anti-β-amyloid, 6E10 (Biolegend, 803014, RRID: AB_2728527), secondary Alexa Fluor™ 488-conjugated goat anti-mouse (1:500, Invitrogen, A-11001, RRID: AB_2534069), mouse anti-β-amyloid (MOAB-2)-Alexa Fluor-647 (1:1000, Novus,: NBP2-13075AF647, RRID:AB_3260696), rabbit anti-amyloid fibrils OC antibody (1:1000, <u>MilliporeSigma</u>, AB2286), secondary Alexa Fluor™ 568-conjugated goat anti-rabbit (1:500, Thermo Fisher,: A-11011, RRID: AB_143157), rabbit anti-GFAP (1:1000, Dako, Z0334), secondary Alexa Fluor™ 568-conjugated goat anti-rabbit (1:500, Thermo Fisher,: A-11011, RRID: AB_143157), rabbit anti-Iba-1 (1:1000, Wako, catalog: 019-19741), secondary Alexa Fluor™ 568-conjugated goat anti-rabbit (1:500, Thermo Fisher, A-11011, RRID: AB_143165), rabbit anti-human amyloid beta (1-40) (1:1000, Immuno-Biological Laboratories, 18580), secondary Alexa Fluor™ 568-conjugated donkey anti-rabbit (1:500, Thermo Fisher, A-10042, RRID: AB_2534017), rabbit anti-human amyloid beta (1-42) (1:1000, Immuno-Biological Laboratories, 18582), secondary Alexa Fluor™ 488-conjugated goat anti-rabbit (1:500, Thermo Fisher, A-11008, RRID: AB_143165), rabbit anti-aquaporin 4 (1:1000, Merck, AB3594), secondary Alexa Fluor™ 568-conjugated goat anti-rabbit (1:500, Thermo Fisher, A-11011,RRID: AB_143157), rabbit anti-CD206 (1:1000, Cell Signaling Technology, mAb #24595), secondary Alexa Fluor™ 568-conjugated goat anti-rabbit (1:500, Thermo Fisher, A-11011,RRID: AB_143157), goat anti Olig2 (1:1000, Biotechne, AF2418), secondary Alexa Fluor™ 647-conjugated donkey anti-goat (1:500, Thermo Fisher, A-21447,RRID:AB_2535864), Alexa Fluor™ 633 Hydrazide for elastin (1:1000, A30634), rabbit anti-alpha smooth muscle actin (1:500, Abcam, ab5694), secondary Alexa Fluor™ 568-conjugated donkey anti-rabbit (1:500, Thermo Fisher, A-10042,RRID: AB_2534017), goat anti CD31(1:800, Bio-Techne, AF3628-SP), secondary Alexa Fluor™ 647-conjugated donkey anti-goat (1:500, Thermo Fisher, A-21447,RRID:AB_2535864), secondary Alexa Fluor™ 633-goat anti-guinea pig IgG (Thermo Fisher, A-21105, RRID:AB_2535757; this secondary antibody was used for 5xFAD mouse IH injections).

#### Imaging

Images were captured with a Zeiss AXIO imager M2 microscope (Germany) and processed with ZEN software (ZEN 2.6, Carl Zeiss, Germany). A 10× objective was used for overview images. Images of the hippocampus, penetrating vessels, astrocytes, and microglia were obtained using 20x, 40x and 63x objectives. Confocal microscopy was performed with a Leica TCS SP8 laser scanning confocal microscope with LAS X software. A 63x objective (HC PL CS2/Oil NA 1.40) was used to image intracellular Aβ (Figure S2) and blood vessels as seen in figure 4b, c, d, e, f, figure 5p, q, r, s. Imaris software (Imaris SingleFull with Clearview version 10.2, Oxford Instruments) was used for 3D reconstruction in Figure S3g, h.

#### Analysis and statistics

All data were analyzed by GraphPad Prism 10.1.2. All normally distributed data sets were assessed with *t* test or One-way ANOVA. All values are displayed as mean ± SD, and N represents the number of mice. We set the statistical significance level at *p* < 0.05. For the IH 6E10 antibody injections, we analyzed the MOAB2 labeling in CA1 neurons, astrocytes, and microglia activation, and MOAB-2 and BSA-647 coverage in the injected and uninjected sides of the hippocampus. For CM 6E10 antibody injections, we analyzed the intracellular MOAB2 in CA1 neurons, astrocytes, and microglia activation, and MOAB-2 and BSA-647 coverage in the whole brain. Intracellular Aβ was analyzed in the dorsal part of the hippocampal field CA1.

## Abbreviations

AD: Alzheimer’s disease
AQP4: aquaporin-4
APP: amyloid precursor protein
ARIA: amyloid-related imaging abnormalities
Aβ: amyloid-β
BACE1: β-site APP-cleaving enzyme 1
BBB: Blood–brain barrier
BSA.: bovine serum albumin
CA1: Cornu Ammonis area 1 (hippocampal region)
CAA: cerebral amyloid angiopathy
CM: cisterna magna
CSF: cerebrospinal fluid
FcγR: Fc gamma receptor
ICV: intracerebroventricular
DS: Down syndrome
FDA: Food and Drug Administration
HS: hypertonic saline
IBA1: ionized calcium-binding adapter molecule 1
IH: intrahippocampal
IP: intraperitoneal
NGS: normal goat serum
NDS: normal donkey serum
NS: normal saline
PBS: phosphate buffered saline
PFA: paraformaldehyde
PS1: presenilin 1
PVM: perivascular macrophage

## Acknowledgements

We acknowledge the technical assistance of Bodil Israelsson and the Multidisciplinary Research Environment on Parkinson’s and Related Diseases MultiPark for their microscopy core facility.

## Author contributions

GHW and GKG designed and supervised the study. GHW, NL, and XNZ performed the experiments. GHW, NL, XNZ and MGR generated and analyzed the statistical data. GHW and XNZ prepared the figures. GHW, NL, TD, AKF and GKG edited the manuscript.

## Data availability

Data are provided within the manuscript or supplementary information files.

## Declarations Ethics approval

All mouse experiments were ethically approved by the Malmo/Lund Ethics Committee on Animal Testing (dnr 5.8.18-13038/2024).

## Consent for publication

All authors have consented for the publication of this manuscript.

