## Supplemental figures for "Aβ Antibodies Target Not Only Amyloid Plaques But Also Distinct Brain Cells And Vessels"

**Supplementary material**
**Supplementary figures and figure legends**

**Fig. S1** Unilateral IH injection of unconjugated Aβ antibody 6E10 and conjugated 6E10 distribution in 5xFAD mice and $APP^{NL\text{-}F}$ mice. (a, b) Unilateral IH injection of unconjugated Aβ antibody 6E10 at 24 and 72 h in 5xFAD mice; scale bar = 1 mm. (c, d) Conjugated Aβ antibody 488-6E10 distributions in astrocytes and microglia after 24 and 72 h of unilateral IH injection in 5xFAD mice; scale bar = 20μm. (e) Secondary antibody distribution after 24 h of unilateral IH injection in 5xFAD mouse; scale bar=1 mm. Conjugated Aβ antibody 488-6E10 distributions in microglia after 24 (f) and 72 h (g) of unilateral IH injection in $APP^{NL\text{-}F}$ mice; scale bar = 20μm.

**Fig. S2** Reduction in intraneuronal Aβ levels of different injection methods of Aβ antibody 488-6E10. (a, d) DAPI staining of CA1 of 4-month-old 5xFAD in 488-6E10 antibody unilateral IH uninjected compared to injection sides. (b, e) MOAB-2 staining for intracellular Aβ of the same area as Figure S2a, d. (c, f) Colocalization of DAPI and MOAB-2 labelling of CA1. (g, j) MOAB-2 labelling of CA1 of 4-month-old $APP^{NL\text{-}G\text{-}F}$ after PBS or 488-6E10 antibody CM injection combined with hypertonic saline IP injection. (h, k) Colocalization of DAPI and MOAB-2 labelling of CA1. (i) Analysis of intracellular Aβ in the uninjected and injected side of CA1 of 5-month-old 5xFAD mice after 24 and 72 h. (l) Analysis of intracellular Aβ in the CA1 after PBS or 488-6E10 antibody CM injection combined with HS IP injections of 4-month-old $APP^{NL\text{-}G\text{-}F}$ mice. N=3 per group. Data are represented as mean ± SD. *$p$<0.05, **$p$<0.01, ***$p$<0.001, scale bar=10 μm.

**Fig. S3** Effect of HS IP injection on CSF influx. (a) 10 μl of BSA-647 was injected into the CM of WT mice. Mice received isotonic saline (normal saline, NS) or hypertonic saline (HS) IP injection at the onset of the CM injection with BSA-647. (b) 10 μl of Aβ antibody 488-6E10 was injected into the CM of $APP^{NL\text{-}G\text{-}F}$ mice. Mice received NS or HS IP injection at the onset of the CM injection with antibody 488-6E10. (c) Representative images from NS and HS groups of WT mice of BSA-647 distribution; scale bar=1 mm. (d) Representative images from NS and HS groups of $APP^{NL\text{-}G\text{-}F}$ mice of Aβ antibody 488-6E10 distribution; scale bar=1 mm. (e) Quantification of BSA-647 distribution in NS and HS groups of WT mice. (f) Quantification of Aβ antibody 488-6E10 distribution in NS and HS groups of $APP^{NL\text{-}G\text{-}F}$ mice. (g, h) 3D reconstruction of Aβ plaques, penetrating vessels in the cortex, and Aβ antibody 488-6E10 showing more antibodies targeting plaques and blood vessel in HS group compared with NS group; scale bar=20 μm. CD31 antibody-blue, amyloid fibrillar oligomer antibody OC-red, Aβ antibody 488-6E10-green. N=3 per group, Data are represented as mean ± SD. *$p$<0.05, **$p$<0.01, ***$p$<0.001, ****$p$<0.0001.

**Fig. S4** Aβ antibody 488-6E10 CM injection combined with HS IP injection reduced plaques and activated microglia and astrocytes in 4-month-old $APP^{NL\text{-}G\text{-}F}$ mice. (a) PBS or Aβ antibody 488-6E10 CM injection combined with HS IP injection of $APP^{NL\text{-}G\text{-}F}$

mice. (b) Images of antibody MOAB2 labelling for Aβ plaques after PBS or 488-6E10 CM injection. Images of IBA-1 (c) and GFAP (d) labelling after PBS or 488-6E10 CM injection. Analysis of MOAB2 (e), IBA-1 (f), and GFAP (g) after PBS or 488-6E10 antibody CM injections; scale bar = 1mm, N=4 per group. Data are represented as mean ± SD. *$p$<0.05, **$p$<0.01.

**Fig. S5** CM injected Aβ antibody 488-6E10 distribution in PVM and oligodendrocytes of 4-month-old $APP^{NL-G-F}$ mice. (a) PVM colocalization with Aβ antibody 488-6E10 in the penetrating vessel of the cortex. (b) Images of antibody CD206 labelling in a penetrating vessel. (c) Image of Aβ antibody 488-6E10 distribution in the penetrating vessels; scale bar=20 µm. (d) Images of perinuclear Aβ antibody 488-6E10 labelling surrounding the nuclear oligodendrocyte-specific antibody Olig2; scale bar=20 µm. (e) Images in higher magnification of Fig. S5d; scale bar = 5 µm. (f) Intracellular Aβ42 in oligodendrocytes; scale bar=5 µm. (g) Colocalization of Aβ antibody 488-6E10 and intracellular Aβ42 surrounding the nuclear oligodendrocyte-specific antibody Olig2; scale bar=5 µm.

**Fig. S6** Distribution of the Aβ antibody 488-6E10 in different brain cells following CM injection in 5-month-old $APP^{NL-F}$ and WT mice. (a) 10 µl of Aβ antibody 488-6E10 was injected into the CM with HS IP injection in $APP^{NL-F}$ mice. (b) The distribution of Aβ antibody 488-6E10 in the brain; scale bar = 1 mm. (c) The distribution of Aβ antibody 488-6E10 in the hippocampus; scale bar = 200 µm. (d, g) Aβ antibody 488-6E10 distribution in CA1 and CA3 neurons; scale bar = 20 µm. (e, h) MOAB2 labelling for intracellular Aβ in CA1 and CA3; scale bar = 20 µm. (f, i) Colocalization of Aβ antibody 488-6E10 and intracellular Aβ, scale bar = 20 µm. Aβ antibody 488-6E10 in PVM (j) of penetrating vessel and microglia (l), surrounding the nuclear oligodendrocyte-specific nuclei antibody Olig2 (k); scale bar = 20 µm, for higher magnification; scale bar = 10µm. (m) Aβ antibody 488-6E10 distribution in the brain of WT mice; scale bar = 1 mm. (n) The distribution of Aβ antibody 488-6E10 in the hippocampus; scale bar = 200 µm. Aβ antibody 488-6E10 in PVMs (o) of penetrating vessels and microglia (p) in the cortex of WT mice; scale bar = 20 µm.

**Fig. S7** Analysis of Aβ antibody 488-6E10 on glymphatic influx and effects on PVM. Analysis of BSA-647 distribution in the hippocampus of 5xFAD mice after unilateral IH injection of Aβ antibody (a), lateral ventricle injection of Aβ antibody 488-6E10 in $APP^{NL-G-F}$ mice (b), and co-injection of Aβ antibody 488-6E10 with BSA-647 in $APP^{NL-G-F}$ mice (c). Image of BSA-647 distribution (d), Aβ antibody distribution (e), and CD206 labelling (f) in the brain after co-injection of Aβ antibody 488-6E10 with BSA-647 in $APP^{NL-G-F}$ mice; scale bar=1 mm. (g) Image of colocalization of BSA-647 and antibody CD206 (PVMs); scale bar=1 mm. (h, i) Colocalization of BSA-647 and CD206 in cortical penetrating vessels and hippocampus; scale bar=100 µm.

Figure S1

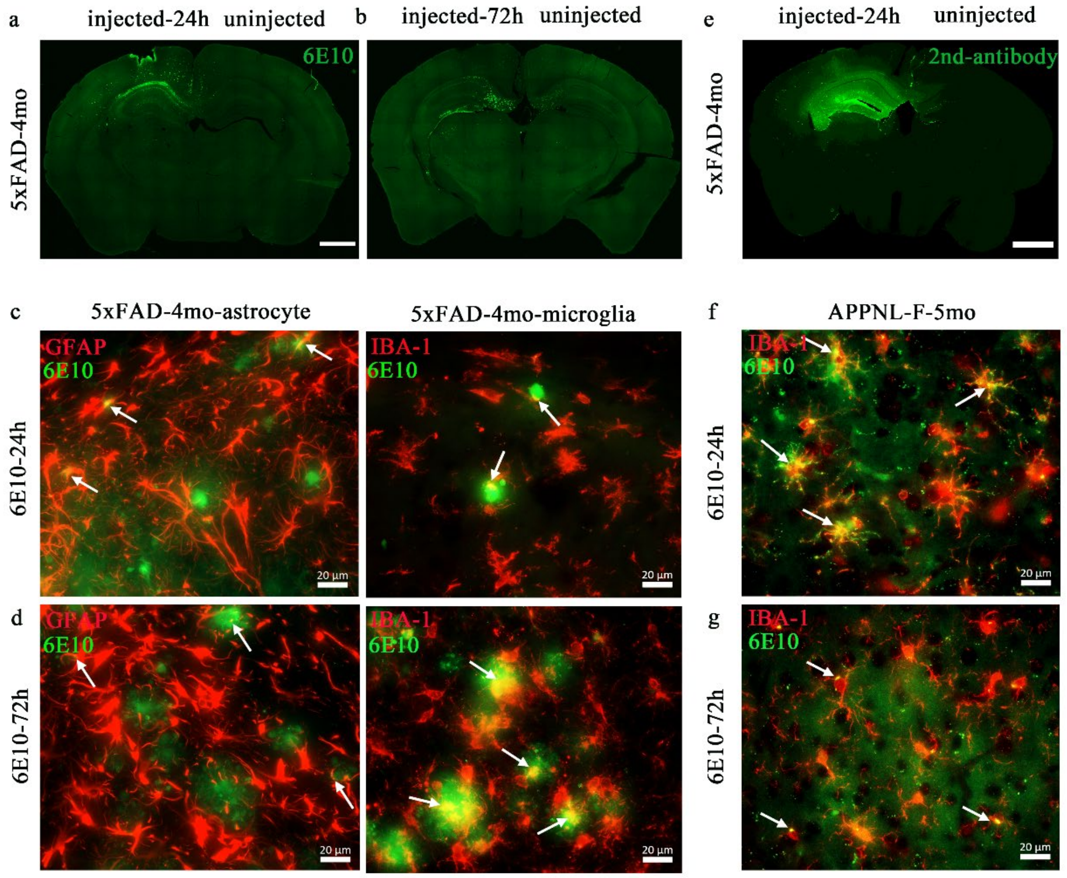
a
injected-24h uninjected
b
injected-72h uninjected
e
injected-24h uninjected
6E10
5xFAD-4mo
2nd-antibody
5xFAD-4mo
c
5xFAD-4mo-astrocyte
5xFAD-4mo-microglia
f
APPNL-F-5mo
GFAP
6E10
IBA-1
6E10
IBA-1
6E10
6E10-24h
6E10-24h
20 µm
d
g
GFAP
6E10
IBA-1
6E10
IBA-1
6E10
6E10-72h
6E10-72h
20 µm

Figure S2

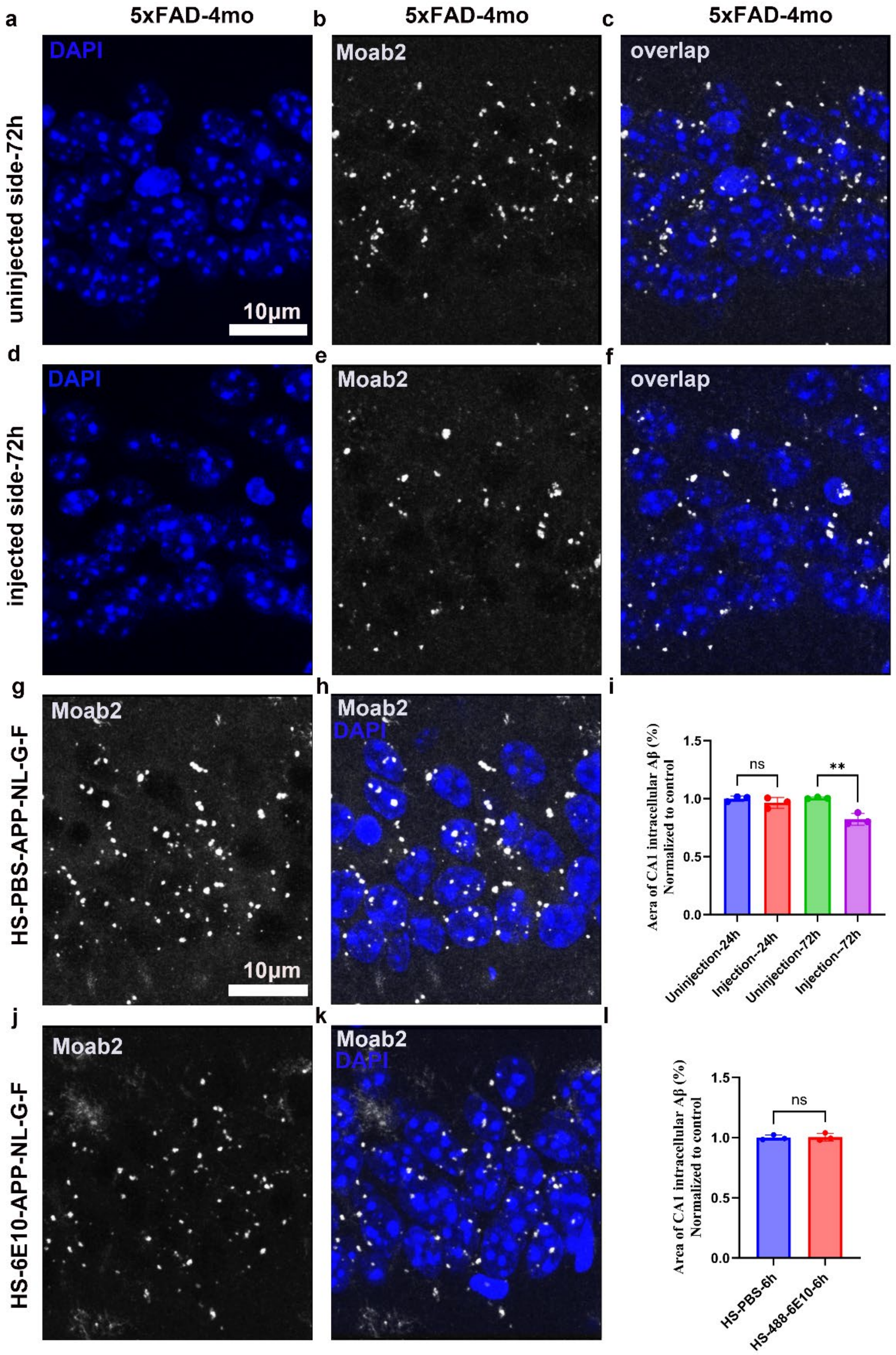

a
b
c
5xFAD-4mo
5xFAD-4mo
5xFAD-4mo
DAPI
Moab2
overlap
uninjected side-72h
10μm
d
e
f
DAPI
Moab2
overlap
injected side-72h
g
h
i
Moab2
Moab2
DAPI
HS-PBS-APP-NL-G-F
10μm
Aera of CA1 intracellular Aβ (%)
Normalized to control
ns
**
1.5
1.0
0.5
0.0
Uninjection-24h
Injection--24h
Uninjection-72h
Injection--72h
j
k
l
Moab2
Moab2
DAPI
HS-6E10-APP-NL-G-F
Area of CA1 intracellular Aβ (%)
Normalized to control
ns
1.5
1.0
0.5
0.0
HS-PBS-6h
HS-488-6E10-6h

Figure S3

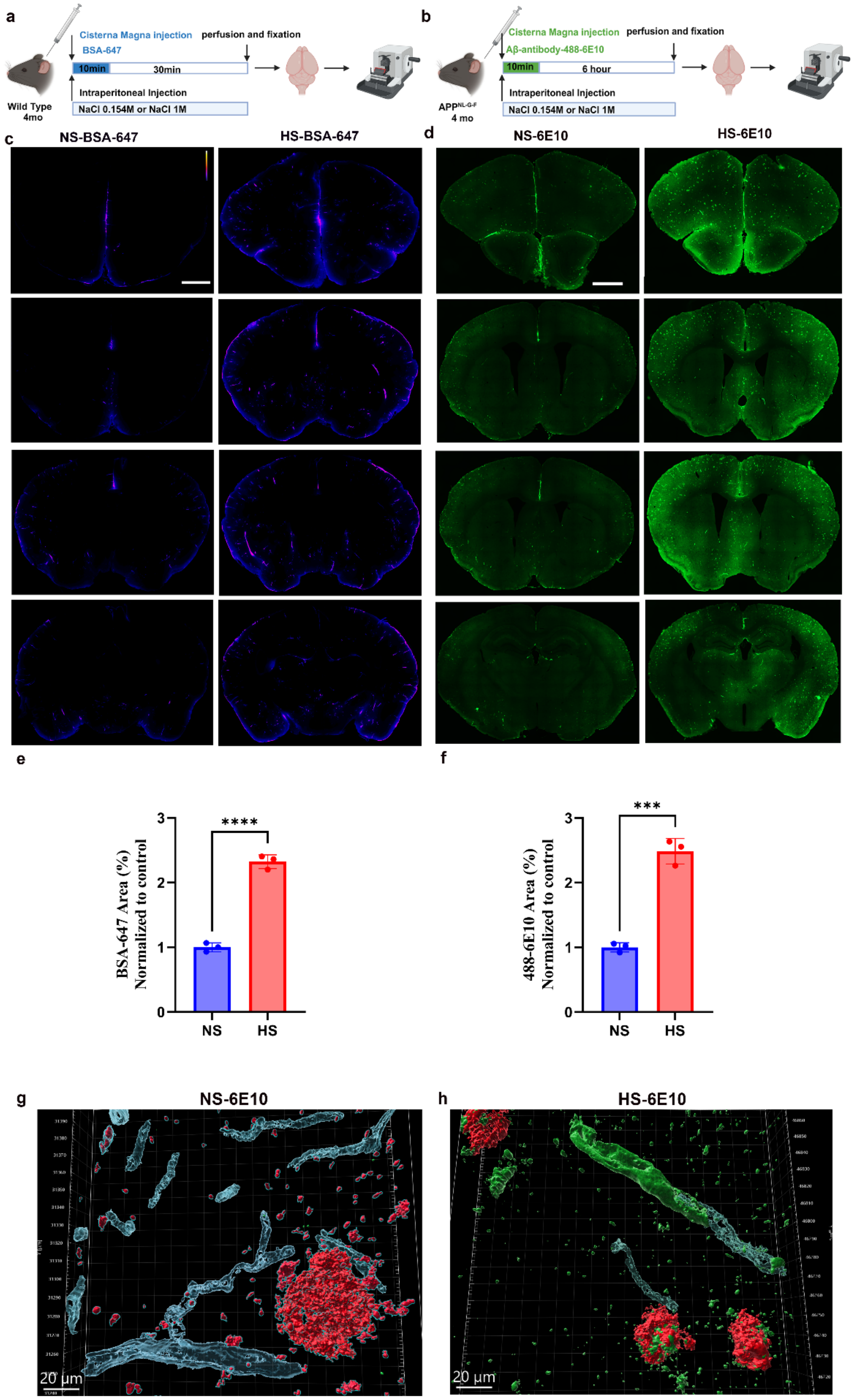
a
Cisterna Magna Injection
BSA-647
perfusion and fixation
10min
30min
Intraperitoneal Injection
NaCl 0.154M or NaCl 1M
Wild Type
4mo
b
Cisterna Magna injection
Aβ-antibody-488-6E10
perfusion and fixation
10min
6 hour
Intraperitoneal Injection
NaCl 0.154M or NaCl 1M
APP NL-G-F
4 mo
c
NS-BSA-647
HS-BSA-647
d
NS-6E10
HS-6E10
e
BSA-647 Area (%)
Normalized to control
****
NS
HS
f
488-6E10 Area (%)
Normalized to control
***
NS
HS
g
NS-6E10
20 μm
h
HS-6E10
20 μm

Figure S4

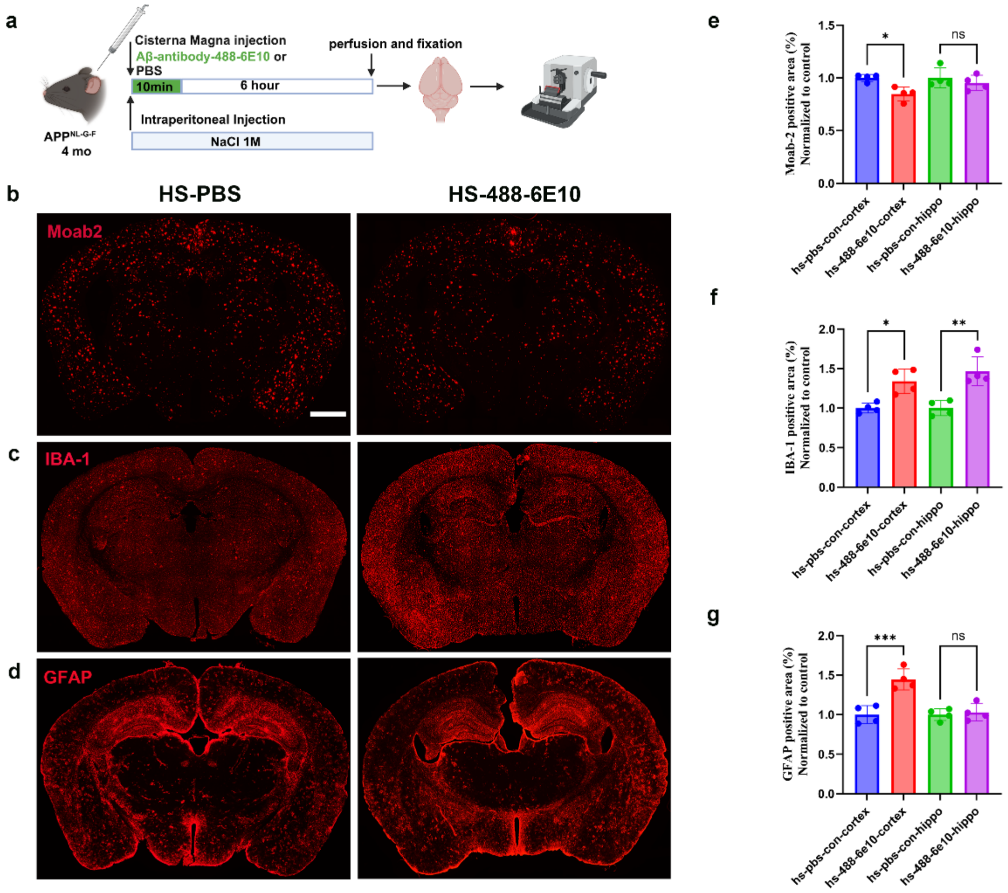
a
Cisterna Magna injection
Aβ-antibody-488-6E10 or
PBS
10min
6 hour
perfusion and fixation
Intraperitoneal Injection
NaCl 1M
APPNL-G-F
4 mo
b
HS-PBS
HS-488-6E10
Moab2
c
IBA-1
d
GFAP
e
Moab-2 positive area (%)
Normalized to control
*
ns
f
IBA-1 positive area (%)
Normalized to control
*
**
g
GFAP positive area (%)
Normalized to control
***
ns
hs-pbs-con-cortex
hs-488-6e10-cortex
hs-pbs-con-hippo
hs-488-6e10-hippo

Figure S5

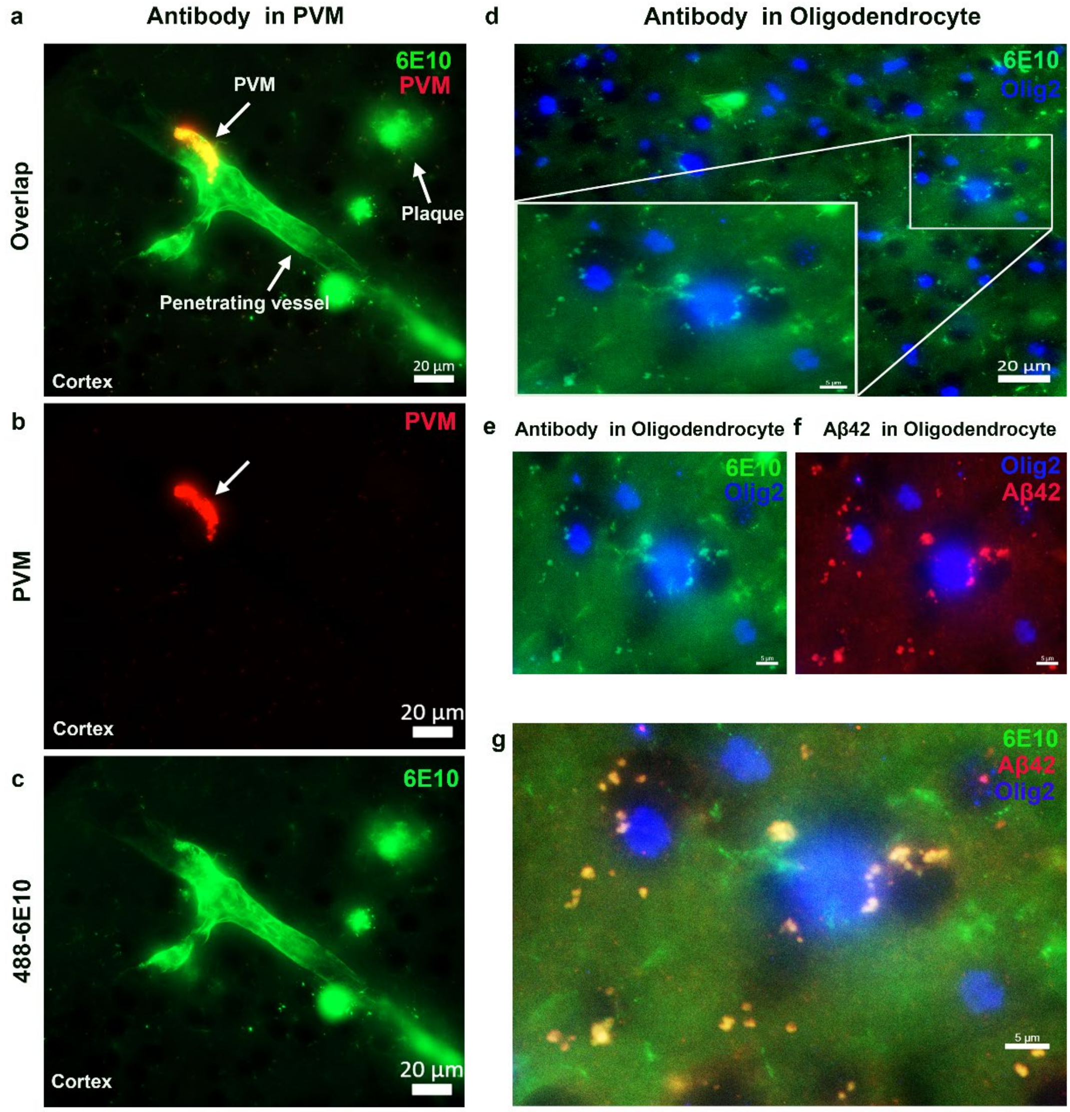

a
Antibody in PVM
Overlap
6E10
PVM
PVM
Plaque
Penetrating vessel
Cortex
20 µm
b
PVM
PVM
Cortex
20 µm
c
488-6E10
6E10
Cortex
20 µm
d
Antibody in Oligodendrocyte
6E10
Olig2
20 µm
e
Antibody in Oligodendrocyte
6E10
Olig2
f
Aβ42 in Oligodendrocyte
Olig2
Aβ42
g
6E10
Aβ42
Olig2
5 µm

Figure S6

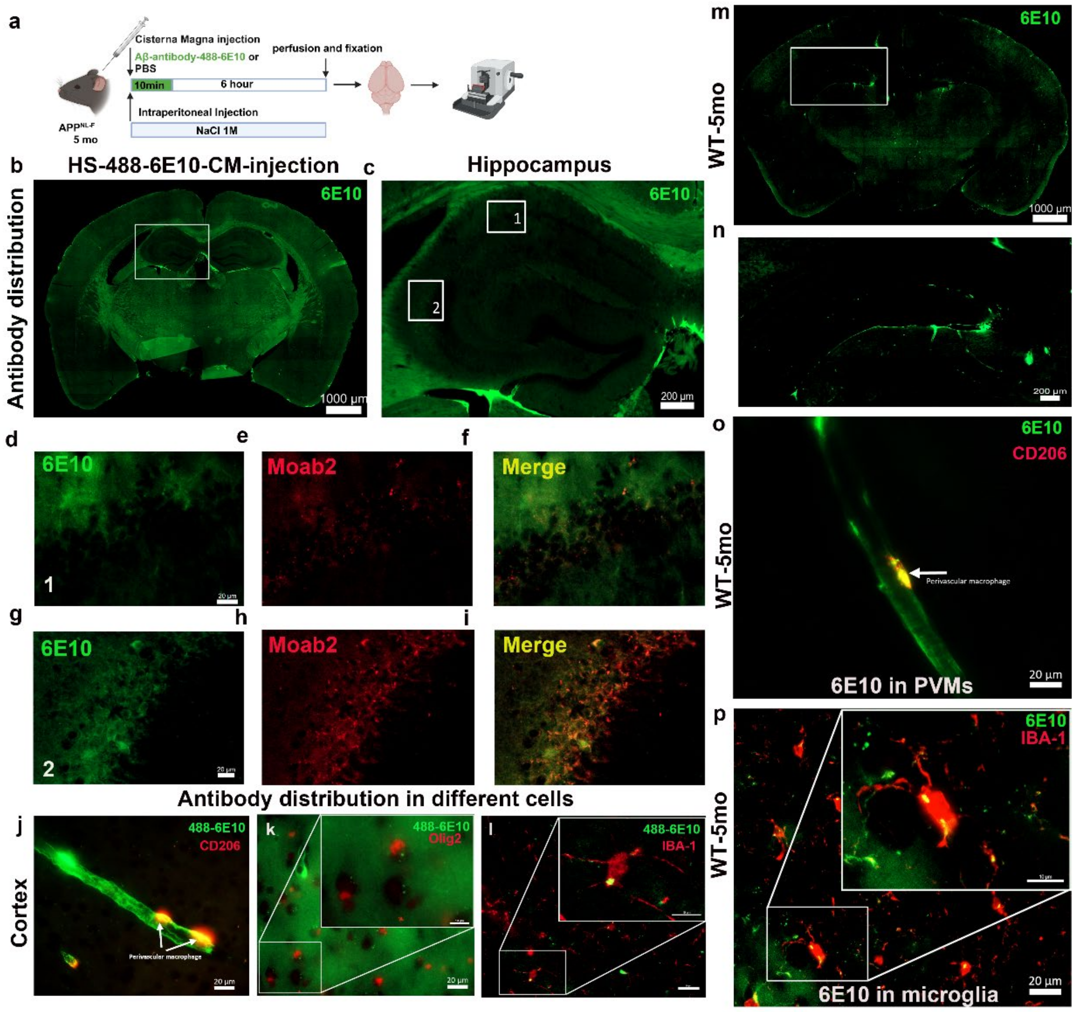

a
Cisterna Magna injection
Aβ-antibody-488-6E10 or PBS
perfusion and fixation
10min
6 hour
Intraperitoneal Injection
NaCl 1M
APP^NL-F
5 mo
b
HS-488-6E10-CM-injection
c
Hippocampus
Antibody distribution
6E10
1000 µm
200 µm
d
e
f
6E10
Moab2
Merge
g
h
i
6E10
Moab2
Merge
Antibody distribution in different cells
j
k
l
Cortex
488-6E10
CD206
Perivascular macrophage
Olig2
IBA-1
20 µm
m
n
o
p
WT-5mo
6E10
CD206
Perivascular macrophage
6E10 in PVMs
IBA-1
6E10 in microglia

Figure S7

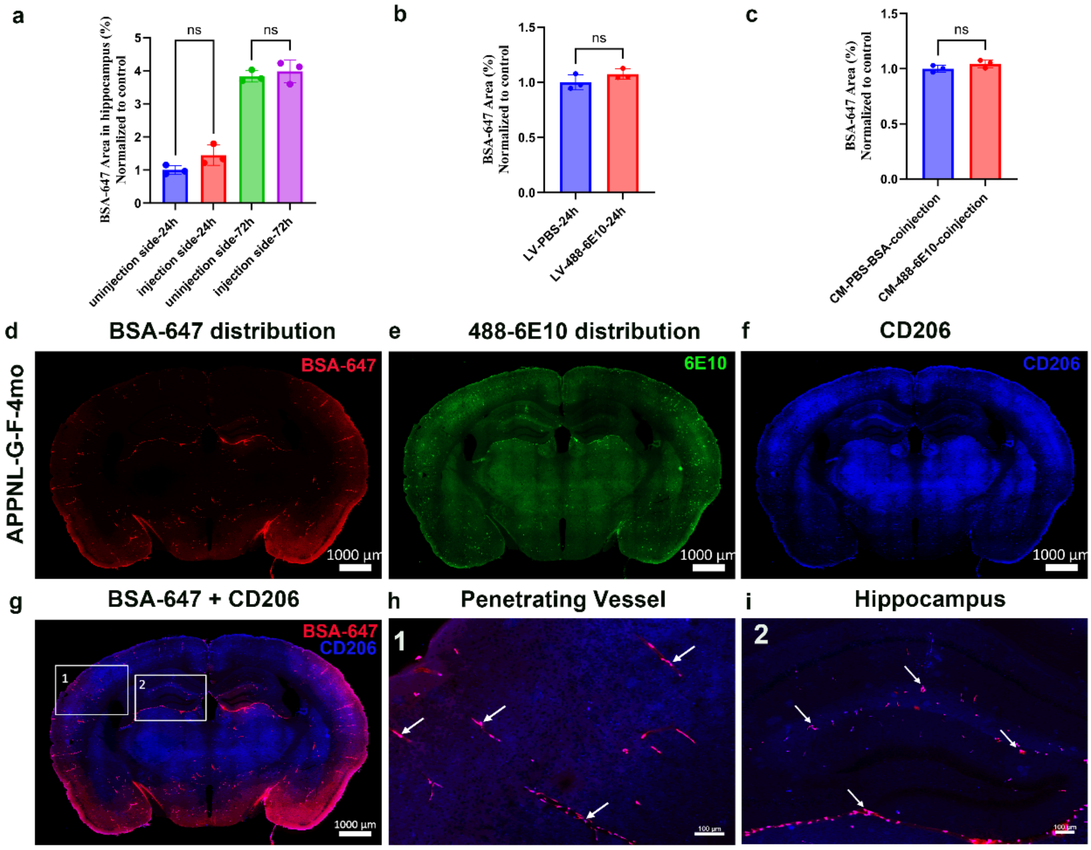
a
b
c
BSA-647 Area in hippocampus (%) Normalized to control
ns
ns
uninjection side-24h
injection side-24h
uninjection side-72h
injection side-72h
BSA-647 Area (%) Normalized to control
LV-PBS-24h
LV-488-6E10-24h
CM-PBS-BSA-coinjection
CM-488-6E10-coinjection
d
BSA-647 distribution
e
488-6E10 distribution
f
CD206
APPNL-G-F-4mo
BSA-647
6E10
CD206
1000 μm
g
BSA-647 + CD206
h
Penetrating Vessel
i
Hippocampus
1
2
100 μm
